# Preventing light-induced toxicity in a new mouse model of rhodopsin sector retinitis pigmentosa

**DOI:** 10.1101/2025.05.14.653774

**Authors:** Rosellina Guarascio, Kaliopi Ziaka, Kwan Hau, Davide Piccolo, Sara Eliza Nieuwenhuis, Adriana Bakoulina, Rowan Asfahani, Monica Aguila, Dimitra Athanasiou, Diana Sefic Svara, Yumei Li, Rui Chen, Michael E Cheetham

**Author notes:** **Correspondence should be addressed to**: Dr Rosellina Guarascio, UCL Institute of Ophthalmology, 11-43 Bath Street, London EC1V 9EL UK.

## Abstract

Retinitis Pigmentosa (RP) is an inherited retinal dystrophy characterized by the progressive loss of rod photoreceptors. Sector RP is a form of RP, where degeneration originates in the inferior retina, mainly influenced by light exposure. Over 200 *RHO* variants are pathogenic and associated with autosomal dominant RP. *RHO^M39R^*is one of the most common *RHO* variants linked to sector RP in the UK. A knock-in (KI) mouse model expressing *Rho^M39R^* was generated and characterized to investigate the mechanisms of degeneration associated with this variant and explore novel therapeutic strategies for rhodopsin sector RP. Under ambient light, *Rho^M39R/+^* KI mice exhibited impaired retinal function by ERG, with some defects in OS ultrastructure, but retained normal outer nuclear layer (ONL) thickness. Repeated exposure to bright light led to photoreceptor loss. In contrast, *Rho^M39R/M39R^* KI mice in ambient light displayed severe retinal dysfunction, ONL thinning, and grossly abnormal OS ultra structure. In homozygous mice, a single bright light exposure significantly reduced ONL thickness within 48 h. The rescue of these models was achieved through reduced light exposure and pharmacological intervention. Rearing in dim red light (red cage condition) restored ERG responses in *Rho^M39R/+^* KI mice and improved ONL thickness in *Rho^M39R/M39R^* KI mice. Transcriptomic analysis in *Rho^M39R/M39R^* KI mice revealed upregulation of Sphingosine 1-P Receptor (S1PR) transcripts. Treatment with the S1PR agonist Fingolimod (FTY720) before bright light exposure significantly reduced degeneration, demonstrating a protective effect in both heterozygous and homozygous models and suggesting potential a therapeutic approach for sector RP patients.

## INTRODUCTION

Retinitis Pigmentosa (RP) is an inherited retinal dystrophy (IRD) that affects 1 in 4,000 people globally. RP leads to the dysfunction and loss of rod photoreceptors followed by the dysfunction and loss of cone photoreceptors (1). In RP, pigment deposits accumulate initially in the peripheral area, while patients experience night blindness and tunnel vision. In later stages of the disease, the degeneration gradually spreads towards the centre of the retina leading to a progressive loss of vision and to legal blindness (1, 2). The progression of the disease can be affected by both genetic and environmental factors, including the exposure of the retina to bright sources of light (3).

*RHO* was the first gene to be associated with RP (4). *RHO*-associated pathology is generally inherited as an autosomal dominant trait (5). *RHO* encodes for rhodopsin, the light-sensitive receptor protein of rod photoreceptor cells that plays a crucial role in the visual cycle and in phototransduction. Structurally, rhodopsin is a G-protein-coupled receptor (GPCR), consisting of an opsin protein bound to 11-*cis*-retinal, a derivative of vitamin A. When light is absorbed, 11-*cis*-retinal isomerizes to all-*trans*-retinal, and rhodopsin undergoes a conformational change that triggers the initiation of the phototransduction pathway essential for vision in scotopic (low-light) conditions (6, 7).

Over 200 *RHO* variants are associated with autosomal dominant forms of RP (adRP). *RHO* variants can be classified according to their biochemical and cellular consequences (5). The best characterized is *RHO^P23H^,* a class 2 mutation that causes misfolding and endoplasmic reticulum (ER) retention of the rhodopsin protein (5, 8, 9). *RHO^P23H^* is also the most frequent *RHO* variant in the US population, with a prevalence of 10-12% among all the adRP cases in the USA (10). A range of transgenic and knock-in (KI) models have been generated to investigate *in vivo* the role of *Rho^P23H^* in the onset and progression of adRP (11-13).

In the UK population, however, *RHO^P23H^* is much less frequently observed, while other less well studied *RHO* variants account for most rhodopsin adRP. One of the most common *RHO* variants in the UK population is *RHO^M39R^. RHO^M39R^* is clinically associated with sector RP, a form of RP where light plays a key role. In sector RP, retinal degeneration initiates in the inferior retina that is more exposed to bright light from above (14). The biochemical and cellular effect of M39R on rhodopsin function was studied using heterologous expression, which showed that it was not retained in the ER and correctly trafficked to the plasma membrane (15). However, rhodopsin M39R was expressed at lower levels, and it showed a higher degree of chemical and thermal instability than WT rhodopsin. Interestingly, rhodopsin M39R caused a faster decay of meta-rhodopsin II and a faster rate of activation of transducin (15). Given these data, *RHO^M39R^* was classified as a class 4 rhodopsin variant, with reduced stability of the rhodopsin protein, but the consequences *in vivo* have not been studied (5).

To investigate the pathogenesis of rhodopsin M39R mediated sector RP, we generated and characterised a *Rho^M39R^* KI mouse model, as a murine model for sector RP. Here we investigated the impact of manipulating light exposure on photoreceptor ultrastructure and survival in *Rho^M39R^*KI mice. Transcriptomic analyses highlighted altered pathways for pharmacological targeting to reveal how molecular interventions can influence the progression of rhodopsin M39R-induced pathology and identify new potential therapeutic approaches.

## RESULTS

### Generation of *Rho^M39R^* mice

The *Rho^M39R^* KI mouse model was generated using homologous-directed repair (HDR) CRISPR/Cas9 technology by electroporating C57BL/6J mouse embryos with a guide RNA (gRNA) and a donor sequence containing nucleotide substitutions for the *Rho* gene (16). A substitution of thymine to guanine at coding position 116 results into an amino acid change from methionine to arginine in position 39, located in the first transmembrane helix (TM) of rhodopsin (Supplementary Table 1, 2). A second substitution was introduced to modify the protospacer adjacent motif (PAM) site and consequently to inhibit Cas9 repeated DNA cleavage (Supplementary Tables 1, 2). Embryos were then reintroduced into a pseudo pregnant mother to generate edited mice. The founder *Rho^M39R/+^* KI mouse was identified by Sanger sequencing. *Rho^M39R/+^* mice were backcrossed with C57BL/6J mice for at least 6 generations to reduce the risk of off-targets effects before performing further experiments (Supplementary Table 3).

### The effect of *Rho^M39R^* expression on photoreceptor survival and function

To investigate the role of rhodopsin M39R on photoreceptor survival and retina activity, *Rho^M39R^* KI mice reared in ambient cyclic light were characterized by optical coherence tomography (OCT), electroretinogram (ERG) (Supplementary Figure 1) and immunohistochemistry (IHC) (Figure 1, Supplementary Figure 2). Live images of the central retina were acquired by OCT to measure the outer nuclear layer (ONL) thickness (Figure 1A, Supplementary Figure 2A). Heterozygous *Rho^M39R/+^* mice had a slightly thinner ONL in the inferior retina at 5 months compared to *Rho^+/+^* mice (Supplementary Figure 2A), but no difference was detected at 3 weeks (Figure 1A, Supplementary Figure 1). In contrast, homozygous *Rho^M39R/M39R^*mice showed a significant reduction of the ONL thickness already by 3 weeks of age (Figure 1A, Supplementary Figure 1).

**Figure 1.**
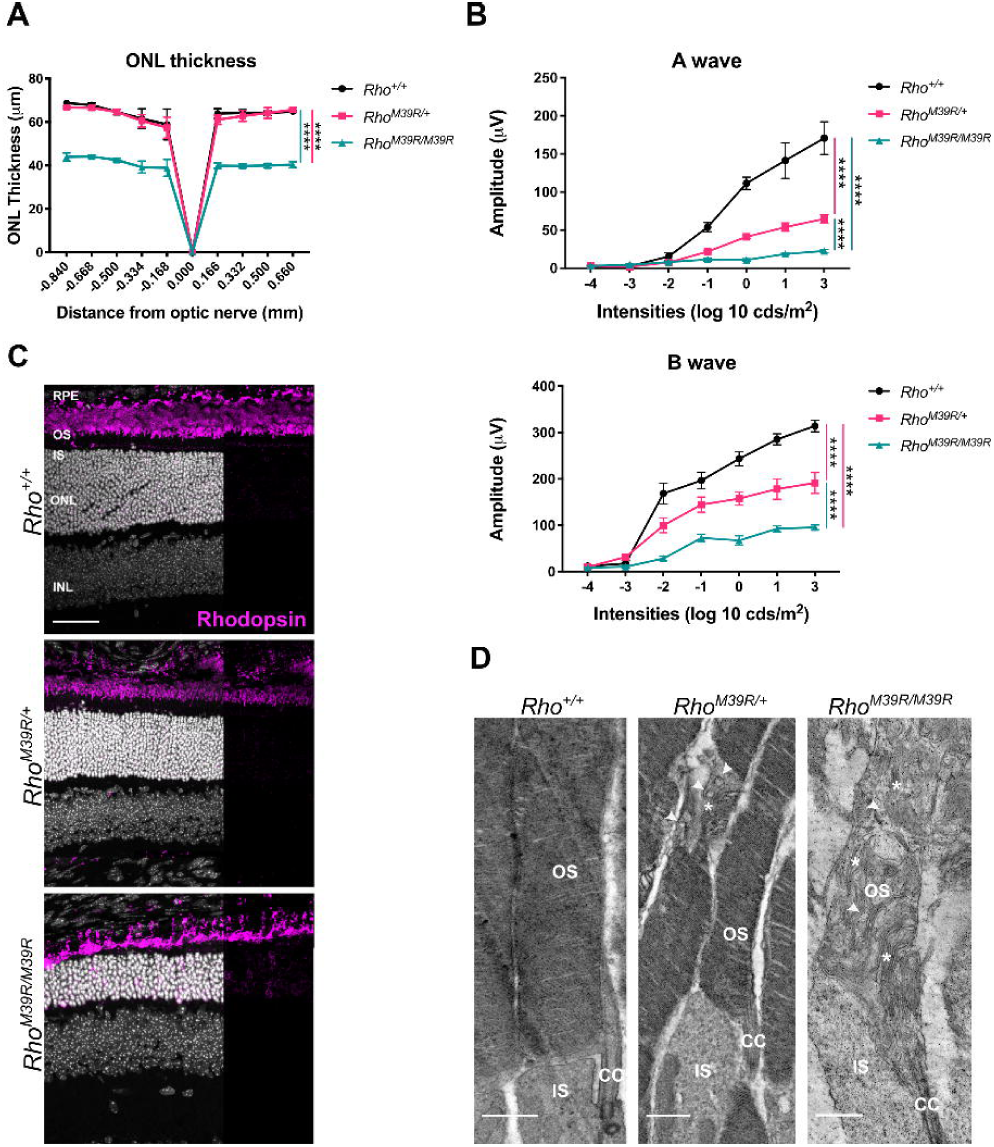
The effect of *Rho^M39R^* expression on photoreceptors. *Rho^M39R/+^* KI, *Rho^M39R/M39R^* KI and *Rho^+/+^* KI mouse models were analysed by OCT, ERG, IHC and TEM at 3 weeks of age. (**A**) The ONL thickness was measured by OCT in the central retina. Mean ± SEM. Two-way ANOVA (**** p<0.0001). N=4. (**B**) The activity of the retina was measured by ERG. The A wave reflects the photoreceptors hyperpolarization in response to different intensities of light and was plotted as positive values. The B wave shows the response of the inner layer of the retina following phototransduction. Mean ± SEM, Two-way ANOVA (**** p<0.0001). N=4 (**C**) Mouse retinae were fixed in 4% PFA for 24 h, incubated in 30% sucrose for 1-2 days, embedded in OCT, cryosectioned and stained with Rhodopsin-4D2 antibody (in magenta). Images were acquired by a Leica Stellaris 8 confocal microscope. Scale bar = 50μm. (**D**) The TEM showed the ultrastructure of the OS in the 3 different models. Stars indicate transversally orientated discs, while arrowheads indicate vesicular structures. Scale bar = 1μm. OS = Outer Segment, CC= connecting cilium, IS = Inner Segment.

Although, *Rho^M39R/+^* mice had no change in photoreceptor survival at 3 weeks, a significant reduction in their light response was detected. Scotopic A and B waves were reduced in *Rho^M39R/+^*compared to *Rho^+/+^* mice retinae (Figure 1B, Supplementary Figure 1B). Furthermore, *Rho^M39R/M39R^* KI mice exhibited a greater deficit in the light response, with scotopic A and B waves values significantly lower than both *Rho^M39R/+^* and *Rho^+/+^*mouse retina (Figure 1B and Supplementary Figure 1B).

### The effect of *Rho^M39R^* on rhodopsin expression and photoreceptor structure

Rhodopsin localised in the outer segment (OS) in both *Rho^M39R/+^*, at 3 weeks and 5 months, and *Rho^M39R/M39R^* models, similar to control retina (Figure 1C, Supplementary Figure 2). By contrast, rhodopsin staining in homozygous mice was also detectable in the rod photoreceptor cell bodies (Figure 1C).

*Rho^P23H^* mice have altered OS discs formation and orientation associated with an impaired ERG (13, 17). Therefore, we investigated the ultrastructure of rod photoreceptor OS in *Rho^M39R^* models by Transmission Electron Microscopy (TEM) at 3 weeks of age (Figure 1D). *Rho^M39R/+^* rod OS were generally comprised of tightly packed discs, similar to *Rho^+/+^* OS. Nevertheless, disorganised areas characterised by transversally oriented discs intercalated with vesicular structures were also observed frequently in the *Rho^M39R/+^* rod OS (Figure 1D). Control wild-type rod OS occasionally had transversally oriented discs, as already described (13, 17); however, the percentage of rod OS with disorganised areas was significantly increased in *Rho^M39R/+^* retinae (Supplementary Figure 3A, B), and OS vesicular structures were only observed in *Rho^M39R/+^*mice (Figure 1D). Furthermore, the OS diameter was significantly smaller by approximately 20% in *Rho ^M39R/+^* compared to control mice. The average diameter was 1.38 ± 0.10μm in *Rho^+/+^* OS, while only 1.13 ± 0.15μm in *Rho^M39R/+^* OS (Supplementary Figure 3C).

Several studies shown that a difference in diameter can be associated with by reduced levels of rhodopsin expression (18, 19). The protein level of rhodopsin in *Rho^M39R/+^* mice was measured by western blot; however, no difference in rhodopsin levels was observed in *Rho ^M39R/+^*mice compared to *Rho^+/+^* mouse retinal lysates (Supplementary Figure 4A, 4B).

In contrast, *Rho^M39R/M39R^* OS were very disrupted with an abnormal orientation and compactness of the discs (Figure 1D). Vesicular structures were also observed. Immunoblot analysis showed that the level of rhodopsin was around 70% lower than in *Rho^+/+^* animals (Supplementary Figure 4A, 4B).

These data show that despite rhodopsin M39R traffic to the OS in *Rho^M39R^* models, photoreceptor function is impaired compared to controls and there is a defect in OS disc organization that is more severe in homozygous mice. In addition, *Rho^M39R/M39R^* had retinal degeneration by 3 weeks of age.

### The role of bright light in *Rho^M39R^* mediated degeneration

*RHO^M39R^* variant is association with sector RP, a form of RP where light is thought to play a critical role (14). Therefore, we hypothesised that light could accelerate dysfunction and degeneration in *Rho^M39R^* mice. In order to assess *Rho^M39R^* photosensitivity, *Rho^M39R/+^* mice were exposed to repeated ERG with a last step of light intensity set up to 30 k lux (Figure 2A). The first ERG was performed on dark-adapted mice at 3 weeks of age and the ERG was repeated two and four weeks later on the same mice. The ONL thickness was measured by OCT as a marker of photoreceptor degeneration. Importantly, the photoreceptor layer was significantly thinner after each round of ERG (Figure 2B). After three ERGs, the ONL thickness of *Rho^M39R/+^* mice reached 15-20μm, equivalent to a few layers of photoreceptors (Figure 2B, 2D). The ERG mediated light exposure significantly reduced the ONL thickness of *Rho^M39R/+^* mice compared to age-matched *Rho^M39R/+^*mice reared in ambient light only at 7 weeks of age (Figure 2C, 2D). Moreover, the repeated light exposure induced a further decline in photoreceptor function in *Rho^M39R/+^* mice, as shown by reduced scotopic A wave amplitude compared to *Rho^M39R/+^* mice reared in ambient light (Figure 2C). There was no change in ONL thickness and photoreceptor activity over 4 weeks in the ambient light only mice. The same paradigm of repeat ERGs on control *Rho^+/+^* mice did not induce degeneration or reduced photoreceptor function (Supplementary Figure 5A, 5B).

**Figure 2.**
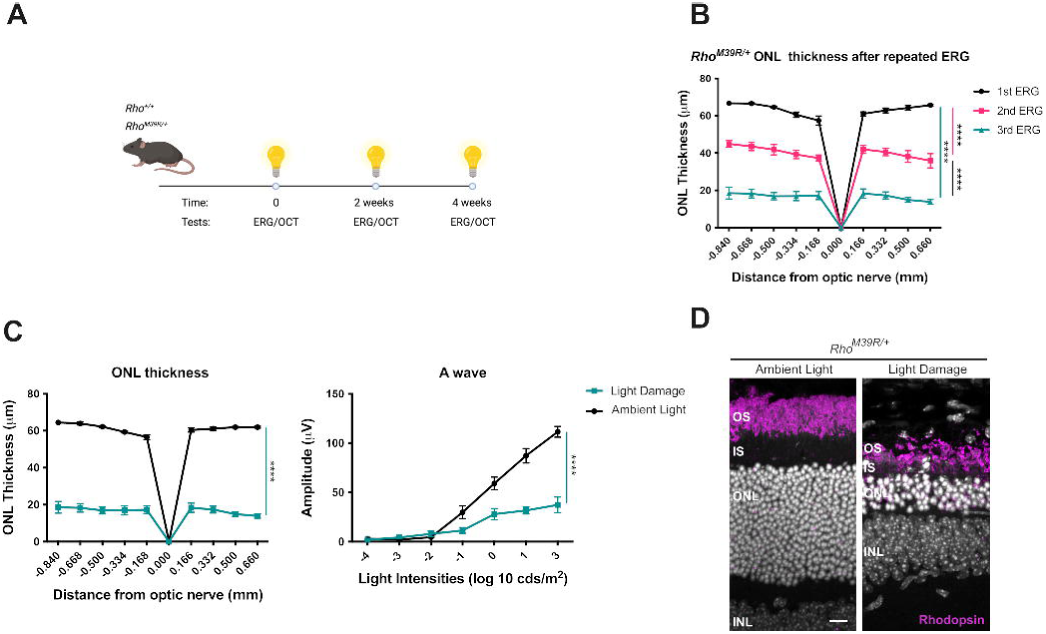
The role of bright light in *Rho^M39R/+^* KI mice. *Rho^M39R/+^* KI mice were exposed to light by ERG at 3 weeks of age. (**A**) Schematic of the light damage assay in *Rho^M39R/+^* KI mice. The animals were dark adapted overnight prior to perform the ERG. The ERG was performed for a total of 3 times, followed by OCT to image the retina and determine the ONL thickness. (**B**) The ONL thickness of the *Rho^M39R/+^*KI mouse retinae was plotted for each round of ERG. Mean ± SEM. Two-way ANOVA (*** p<0.001). N=4 (**C**) *Rho^M39R/+^* KI mice from the light damage assay were compared to animals reared in ambient light to show the difference in ONL thickness and photoreceptors activity (A wave). Mean ± SEM. Two-way ANOVA (**** p<0.0001). N=4. (**D**) IHC of the retina of *Rho^M39R/+^* KI mouse after light damage or reared in ambient light. The cryosections were stained with rhodospin-4D2 antibody (in magenta). Scale bar=10μm.

The same repeat ERG light damage paradigm was also performed on homozygous *Rho^M39R/M39R^* mice. Initially, the first ERG was performed on mice at 3 weeks of age and repeated after a week. However, a week later, the photoreceptor layer was greatly reduced when imaged by OCT (data not shown). Therefore, a single ERG was performed, and the retinae were collected at 3, 6 and 24 h after the bright light exposure (Figure 3A). No changes in the ONL thickness were observed by IHC 3 or 6 hours after bright light exposure. There was in increase in TUNEL reactivity after 3 or 6 h, but this was not statistically significant (Figure 3B-D). Whereas ultrastructural investigation by TEM showed that swollen mitochondria with disrupted cristae were present in the IS of photoreceptors from 3 h after bright light exposure. The alteration in mitochondrial morphology was exacerbated at 6 h (Supplementary Figure 6). Shorter OS full of vesicular structures were also observed in *Rho^M39R/M39R^* mice retina after light exposure (Supplementary Figure 6). Furthermore, by 24 h after bright light ONL thickness had significantly reduced, together with a significant increase in TUNEL reactivity (Figure 3B-D). These data confirm that *Rho^M39R^* mice are sensitive to light to the extent that a short exposure to bright source of light can induce photoreceptor cell death.

**Figure 3.**
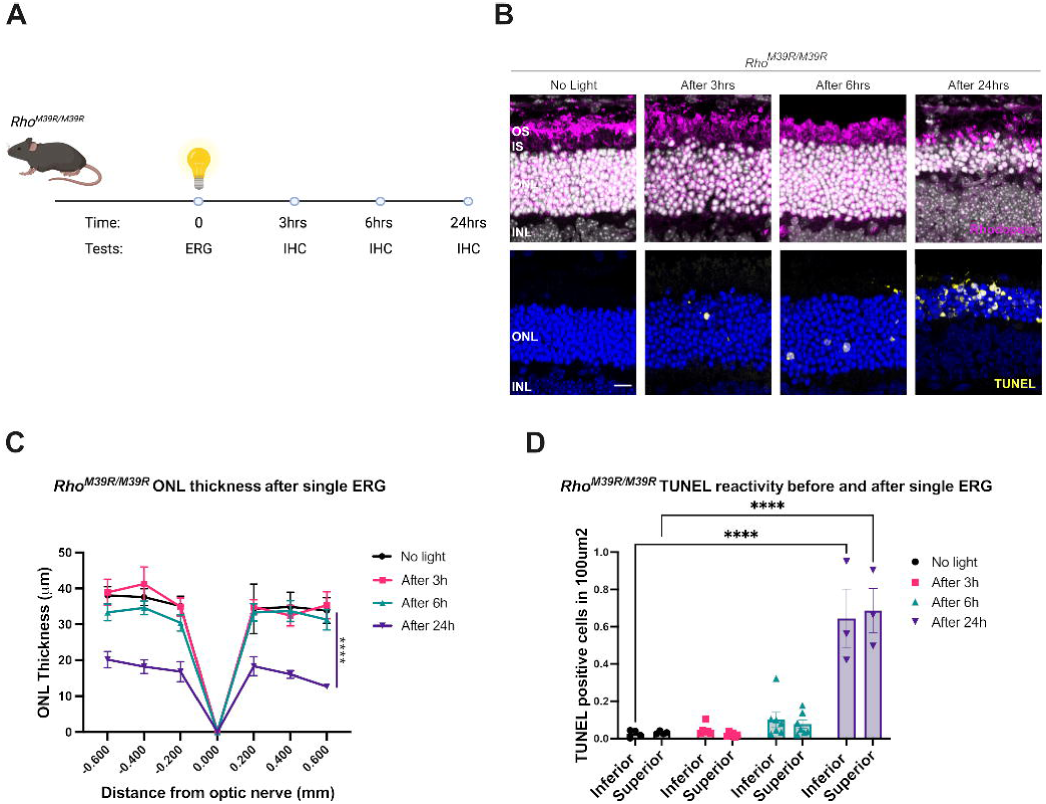
The role of bright light in *Rho^M39R/M39R^* KI mice. *Rho^M39R/M39R^* KI mice were exposed to light by ERG at 3 weeks of age. (**A**) Schematic of the light damage assay in *Rho^M39R/M39R^*KI mice. Mice were dark adapted overnight prior to perform the ERG. The ERG was performed only once. The animals were culled at different time points (3, 6, 24hr after light) and one eye was collected for IHC. (**B**) Mouse retinae were fixed overnight in 4% PFA, incubated in 30% sucrose for 1-2 days, embedded in OCT (embedding matrix), cryosectioned and stained with Rhodopsin-1D4 antibody (in magenta) or used for a fluorochrome-based TUNEL assay (in yellow). Images were acquired using a Zeiss LSM700 confocal microscope. Scale bar=10μm (**C**) Images of the central retina, including the optic nerve, were acquired with an EVOS FL auto 2 microscope and used to measure the ONL thickness at specific distances from the optic nerve comparable to the OCT analysis. (**D**) The number of TUNEL positive cells in the ONL was counted using images of the central retina acquired with an EVOS FL auto 2 microscope. Mean ± SEM. Two-way ANOVA (**** p<0.0001). N=4/7.

### Dim red light protection against *Rho^M39R^*

In order to test the role of ambient light in photoreceptor dysfunction and degeneration, *Rho^M39R^* mice were reared in red cages. Rhodopsin is not activated by red light and rodents do not express an L cone opsin, therefore rodents do not show any visual activity under diminished red light and they can be reared in red cages as if they were living in absence of light (20). *Rho^M39R/+^* mice were reared in dim red light from birth until 3 weeks of age and then characterized. The ONL of *Rho^M39R/+^*mice reared in dim red light was significantly thicker than the ONL thickness of *Rho^M39R/+^*mice or control mice reared in ambient light (Figure 4A). In addition, there was a significant improvement of the scotopic A and B wave amplitude in the *Rho^M39R/+^*animals reared in dim red light compared to *Rho^M39R/+^* mice reared in ambient light. The amplitude of both A and B wave was fully rescued and comparable to the ERG of *Rho^+/+^* mice (Figure 4B). Investigation by TEM revealed that the OS ultrastructure of red cage reared *Rho^M39R/+^* mice was improved, with less frequent breaks observed along the OS compared to animals reared in ambient light (Figure 4C, D).

**Figure 4.**
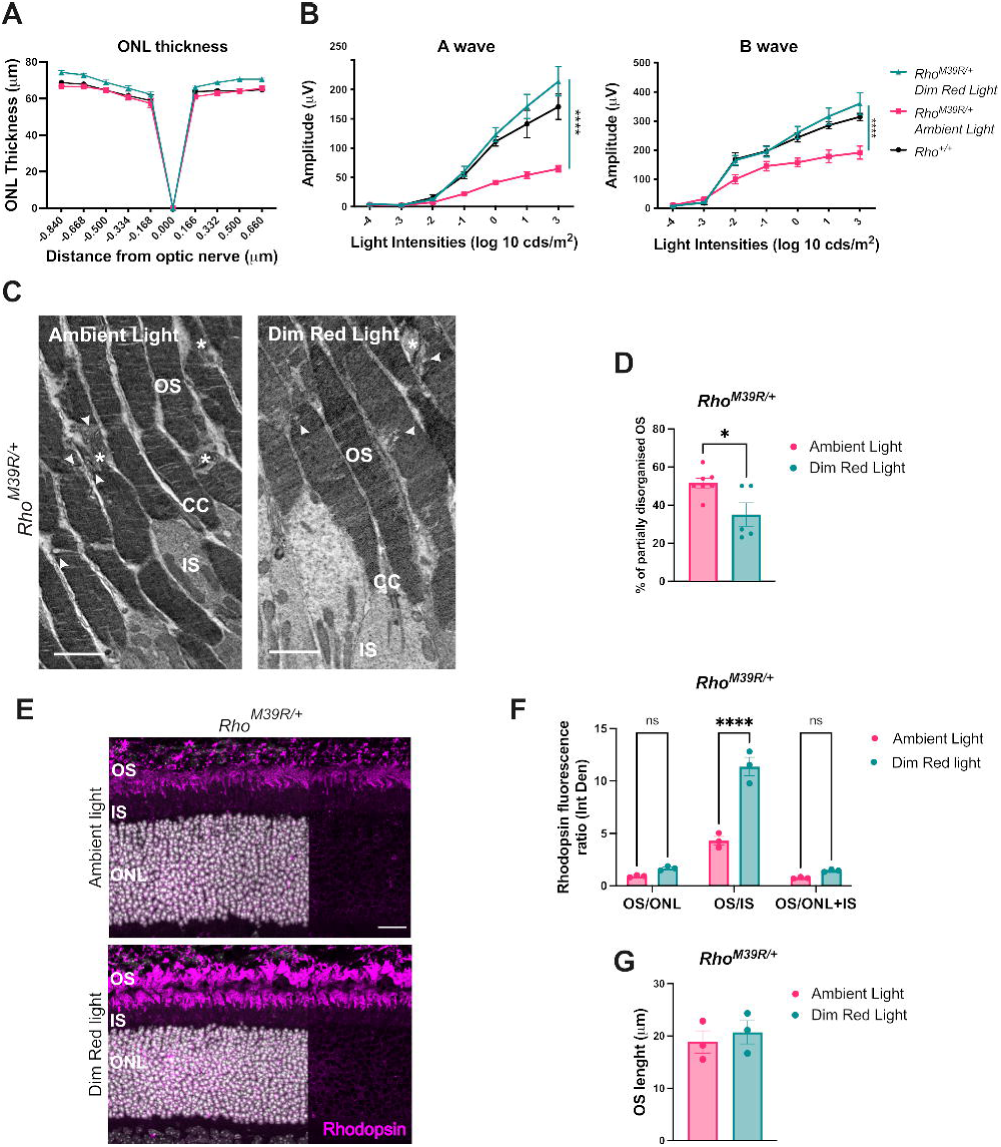
The role of dim red light in *Rho^M39R/+^* KI mice. *Rho^M39R/+^* KI mice were reared in red cages under dim red light until 3 weeks of age. (**A**) The ONL thickness of *Rho^M39R/+^* KI (N=7) reared in dim red light was compared to the ONL thickness of respectively *Rho^+/+^* and *Rho^M39R/+^*KI mice reared in ambient light. The measurements were taken by OCT. (**B**) Scotopic A and B waves were recorded for the same set of animals: *Rho^M39R/+^* KI in dim red light and ambient light; and *Rho^+/+^* KI mice reared in ambient light. Mean ± SEM. Two-way ANOVA (**** p<0.0001). (**C**) The TEM images showed the ultrastructure of *Rho^M39R/+^* KI mice reared in dim red light and ambient light. Stars identify transversally orientated discs, while arrowheads indicate vesicular structures. Scale bar=2μm. OS = Outer Segment, CC= connecting cilium, IS = Inner Segment. (**D**) The % of partially disorganised RO (with transversally orientated discs and/or vesicular structures) was plotted. Mean ± SEM. Mann-Whitney test (*p<0.05). 5-7 images from 2 independent animals. (**E**) Eyes from mice in red cages were enucleated after ERG and OCT and fixed in 4% PFA. Eye cryosections were stained with anti-rhodopin-4D2 antibody (in magenta) Scale bar=20μm. (**F**) The integrated density of rhodopsin signal was measured in the different layers of the retina by using Fuji and plotted as a ratio. Mean ± SEM. Two-way ANOVA (*p<0.05, ** p<0.01). N=3. (**G**) The length of the OS was measured by using Fuji. Mean ± SEM. Unpaired t-test. OS = Outer Segment, CC = connecting cilium, IS = Inner Segment, ONL = Outer Nuclear Layer, INL = Inner Nuclear Layer.

Rhodopsin localization was also investigated in the retina of *Rho^M39R/+^* mice reared in ambient light and dim red light. The relative fluorescence intensity of rhodopsin staining in OS was measured in comparison to other compartments (Figure 4E-G). Interestingly, the rhodopsin OS/IS ratio was significantly higher in the mice reared in dim red light compared to mice reared in ambient light (Figure 4F). No change in OS length was observed (Figure 4G). In summary, ambient light can affect rod photoreceptor activity, rhodopsin localization and OS structure in *Rho^M39R/+^* mice.

Rearing in red cages was also tested in *Rho^M39R/M39R^* mice. Importantly, a significant increase in the ONL thickness was observed in *Rho^M39R/M39R^* mice reared in red cages compared to ambient light reared *Rho^M39R/M39R^* mice (Figure 5A). Nevertheless, ERG analysis did not show any improved retinal function (Figure 5B), even though by TEM, we observed an improvement in the RO ultrastructure. *Rho^M39R/M39R^* mice reared in dim red light showed short OS composed of tight and evenly arranged discs (Figure 5C).

**Figure 5.**
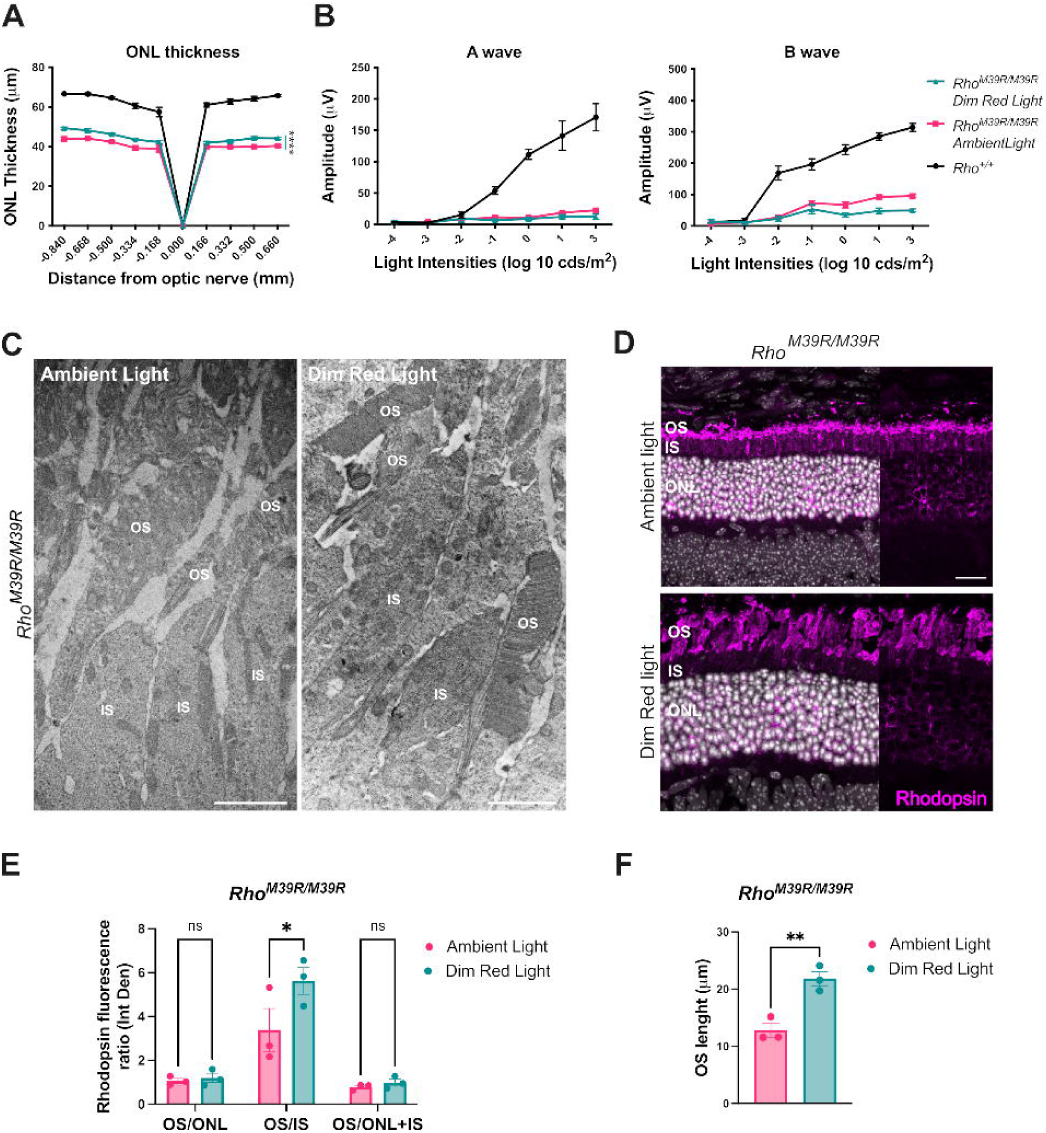
The role of dim red light in *Rho^M39R/M39R^* KI mice. *Rho^M39R/M39R^* KI mice were reared in red cages under dim red light until 3 weeks of age. (**A**) The ONL thickness of *Rho^M39R/M39R^* KI mice (N=5) reared in dim red light was compared to the ONL thickness of mice reared in ambient light. The ONL thickness of *Rho^+/+^* mice reared in ambient light was also plotted. The measurements were taken by OCT. (**B**) Scotopic A and B wave of *Rho^M39R/M39R^* KI in dim red light and ambient light and *Rho^+/+^* mice reared in ambient light was also reported. Mean ± SEM. Two-way ANOVA (**** p<0.0001). (**C**) The ultrastructure of *Rho^M39R/M39R^*KI mice reared in dim red light and ambient light was analysed by TEM. Scale bar=2μm. OS = Outer Segment, IS = Inner Segment. (**D**) Eyes from mice in red cages were enucleated after ERG and OCT and fixed in 4% PFA. Eye cryosections were stained with anti-rhodopin-4D2 antibody (in magenta). (**E**) The integrated density of rhodopsin signal was measured in the different layers of the retina by using Fuji and plotted as a ratio. Mean ± SEM. Two-way ANOVA (*p<0.05, **p<0.01). N=3. (**F**) The length of the OS was measured by using Fuji. Mean ± SEM. Unpaired t-test (**p<0.01). OS = Outer Segment, CC = connecting cilium, IS = Inner Segment, ONL = Outer Nuclear Layer, INL = Inner Nuclear Layer

*Rho^M39R/M39R^* mice raised in ambient light had some rhodopsin immunoreactivity in the IS and in the ONL (Figure 5D). The retention of rhodopsin M39R in the IS was partially rescued in the retina of mice reared in dim red light as shown by a significant improvement in the OS/IS ratio (Figure 5E). Moreover, *Rho^M39R/M39R^* reared in dim red light displayed a longer OS than mice reared in ambient light (Figure 5F). Therefore, ambient light appears to contribute to photoreceptor degeneration in *Rho^M39R/M39R^* mice.

### Altered sphingosine pathway transcripts in *Rho^M39R/M39R^* KI mice

To investigate the transcriptomic changes associated with retinal degeneration driven by *Rho^M39R^*, we performed bulk RNA-sequencing (RNA-seq) and compared gene expression of 3 weeks old *Rho^+/+^*and *Rho^M39R/M39R^* mice. The top differentially expressed genes (DEGs) included genes involved in the inflammatory response and complement activation (Figure 6A). Gene Set Enrichment Analysis (GSEA) using WebGestalt was performed to interpret the upregulated and downregulated DEGs and explore biological processes and pathways involved in retinal degeneration. This confirmed a significant upregulation of inflammatory pathways and a significant downregulation of the phototransduction cascade (Supplementary Figure 7A, B). Interestingly, also genes involved in the sphingolipid signalling pathways were upregulated (Figure 6A, Supplementary Figure 7C). *S1pr3*, *S1pr2* and *Tnfrsf1a* were significantly upregulated in *Rho^M39R/M39R^*compared to control retinae (Figure 6A, Supplementary Figure 7C).

**Figure 6.**
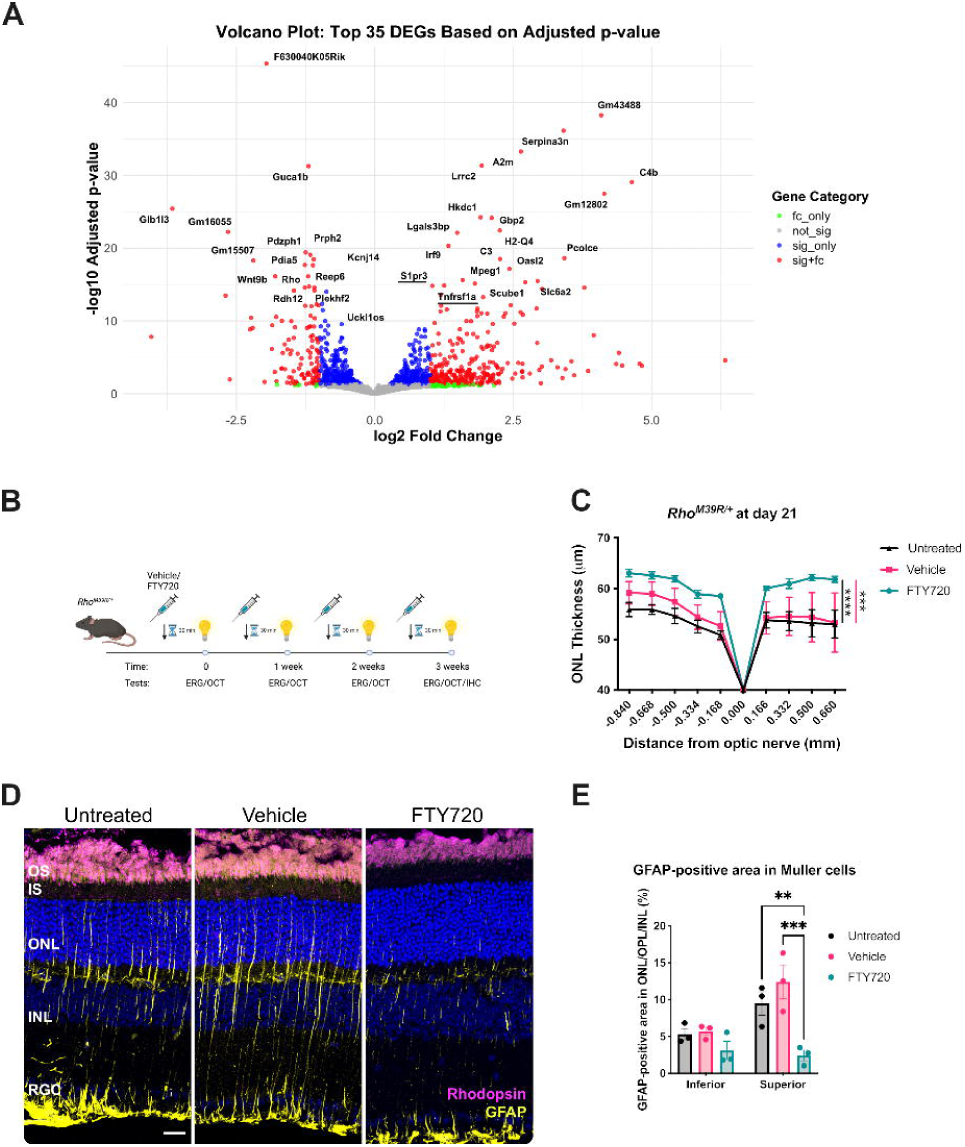
Modulating S1P signaling pathway as a pharmacological treatment for *Rho^M39R/+^* light-induced degeneration. (**A**) Transcriptomic analysis by bulk RNA-seq of *Rho^M39R/M39R^* KI retinae compared to control mice. The volcano plot shows the top 35 differentially expressed genes by adjusted p-value. S1P signalling pathway genes are underlined in red. fc_only = only fold change; not_sign = not significant; sig_only = only significant; sig+fc = significant plus fold change. (**B**) *Rho^M39R/+^* and *Rho^M39R/M39R^* KI mice were treated with FTY720 at 4 weeks of age. Schematic of the treatment in *Rho^M39R/+^* KI mice. The animals were dark adapted overnight and intraperitoneally injected with 10mg/kg of FTY720 or vehicle (saline solution) 30min before performing the light damage assay. The light damage consisted in performing an ERG every week for 4 times in total. Each time, the mice were preinjected with FTY720 or vehicle. (**C**) The ONL thickness was measured at day 21, after 4 rounds of ERG. FTY720-treated mice were compared to vehicle-treated and untreated mice. Mean ± SEM. Two-way ANOVA (**** p<0.0001, *** p<0.001). N= 5/7 (**D**) At the end of the light damage experiments, mice were culled, and the eyes were enucleated to perform further histological investigations. Rodents eyes were fixed in 4% PFA, incubated in 30% sucrose for 1-2 days, embedded in OCT (embedding matrix), cryosectioned and stained with DAPI. IHC of untreated, vehicle or FTY70 treated *Rho^M39R/+^* superior retina after light damage. The cryosections were stained with rhodospin-4D2 (in magenta) and anti-GFAP (in yellow). Scale bar = 20μm. (**E**) The % area occupied by GFAP-positive signal relative to the ONL, OPL and INL was measured after thresholding GFAP staining in untreated and treated animals. Analysis performed in Fiji. Mean ± SEM. Two-way ANOVA (** p<0.001, *** p<0.001). N= 3.

Sphingosine-1-phosphate (S1P) is a sphingolipid that act as a lipid mediator and can modulate several cellular processes including mechanisms of survival and proliferation (21). High levels of S1P and of S1P receptors (*S1pr*) have been associated with light damage in other animal models (22-24). Reduction of phototransduction protein gene expression could be related to the retinal degeneration occurring in *Rho^M39R/M39R^* mice; however, upregulation of sphingolipid signalling pathways suggested they could be involved in the rhodopsin M39R-mediated mechanisms of cell death.

### S1P receptor as a pharmacological target for Rhodopsin M39R light-induced degeneration

Sphingolipid signalling pathways can be pharmacologically targeted. Several small molecules targeting the S1P receptors, such as Fingolimod/FTY720, are effective in preserving cell viability in pathological conditions (25, 26). To investigate if *S1pr2* and *S1pr3* play a critical role in light-induced photoreceptor degeneration in our model, we treated both *Rho^M39R/+^* and *Rho^M39R/M39R^* mice with FTY720 to modulate S1P receptors. Previous reports on light-induced retinal stress treated animals just before the light exposure (22, 27). Therefore, we injected the mice with 10 mg of FTY720/kg body 30min prior to induction of light damage with ERG. *Rho^M39R/+^* mice were injected before each ERG and each ERG was repeated every week for 4 rounds (until day 21) starting with mice at 4 weeks of age (Figure 6B). The last ERG was performed to investigate potential changes in retina activity following the treatment (Supplementary Figure 8C). *Rho^M39R/+^* mice were treated with FTY720, vehicle or left untreated. The ONL thickness was measured at each time point in both treated and untreated mice to monitor the degree of degeneration (Supplementary Figure 8A, 8B). After repeated ERG, the mice treated with FTY720 had a thicker ONL than the untreated and vehicle-treated *Rho^M39R/+^*mice (Figure 6C); however no changes were detected in ERG responses (Supplementary Figure 8C). The ONL thickness in FTY720-treated mice decreased a little compared to day 0, but the rate of degeneration was much slower than untreated and vehicle-treated animals (Supplementary Figure 8B). The degeneration in untreated and vehicle-treated retinae was observed predominantly in the superior retina. Measurement of the number of photoreceptors in fixed and cryosectioned *Rho^M39R/+^* retinae confirmed the improved photoreceptor survival following treatment (Supplementary Figure 8D). In addition, in untreated and vehicle-treated animals at day 21 Müller cells were activated and expressed GFAP; whereas, this reactivity was reduced in FTY720-treated retinae (Figure 6D, E). The GFAP-positive area occupied by reactive Müller cells in the ONL, the outer plexiform layer (OPL) and inner nuclei layer (INL) was significantly decreased in the superior retina of FTY720-treated retinae compared to untreated and vehicle-treated retinae (Figure 6E). A smaller reduction was observed also in the inferior retina, even if not statistically significant (Figure 6E). *Rho^+/+^* mice were also injected with FTY720 or vehicle to confirm the drug safety. No changes in the ONL thickness or retina activity were observed (Supplementary Figure 9A, 9B).

4 week old *Rho^M39R/M39R^* mice were injected with FTY720 before a single ERG, while OCT measurement was performed at day 0 and after 2 days (Figure 7A). FTY720-treated mice had less retinal degeneration than untreated and vehicle-injected mice (Figure 7B). Similar to the *Rho^M39R/+^*mice, the administration of FTY720 did not completely prevent retinal degeneration (Supplementary Figure 10B). Nevertheless, the FTY720 treatment prior to light exposure significantly reduced the amount of photoreceptor loss (Figure 7C, D). However, Müller glia activation did not differ in FTY720-treated retina compared to untreated and vehicle-treated (Supplementary Figure 10C, D).

**Figure 7.**
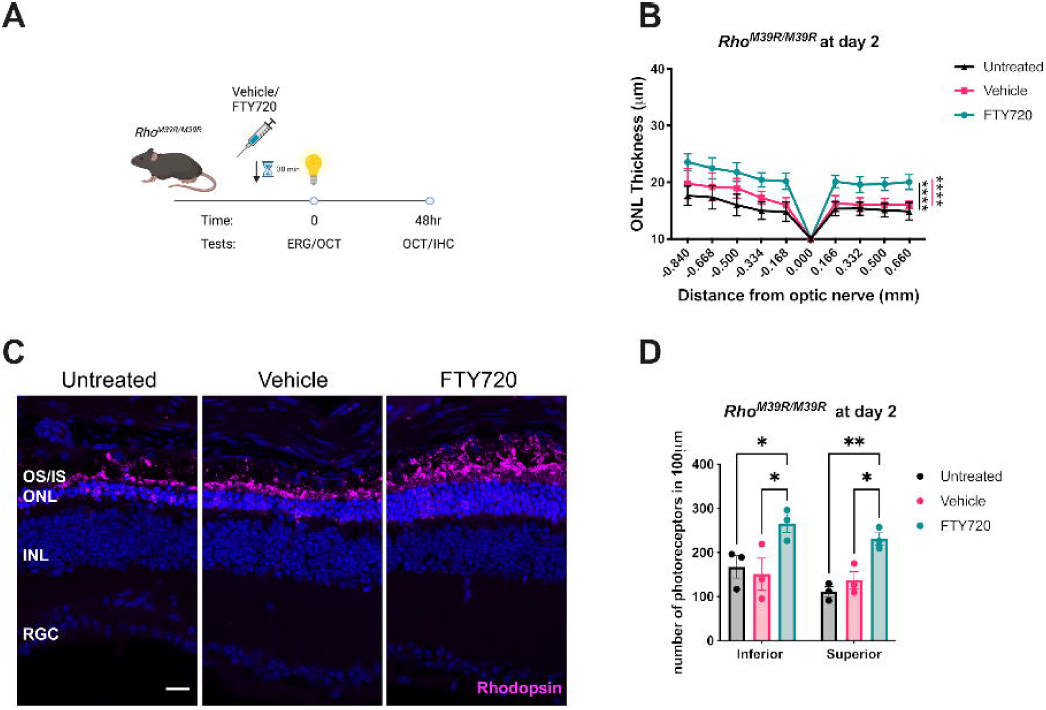
Modulating S1P signaling pathway as a pharmacological treatment for *Rho^M39R/M39R^* light-induced degeneration. (**A**) Schematic of the FTY20 treatment in *Rho^M39R/M39R^* KI mice. Mice were dark-adapted and injected with FTY720 (10 mg/kg) 30 min prior to the light damage assay. The ERG was performed once to induce a faster degeneration. The OCT was performed at both day 0 and day 2, after 48 h from the single ERG. (**B**) The ONL thickness was measured at day 2, after a single ERG. FTY720-treated *Rho^M39R/M39R^* KI mice were compared to vehicle-treated and untreated mice. Mean ± SEM. Two-way ANOVA (**** p<0.0001, *** p<0.001). N=5/6 (**C)** IHC of untreated, vehicle or FTY70 treated *Rho^M39R/M39R^*superior retina after light damage. The cryosections were stained with rhodospin-4D2 (in magenta). Scale bar=20μm. (**D**) The number of photoreceptors in the ONL was measured at 200 to 400 μm from the optic nerve in the inferior and superior retina. The analysis was performed on images of the central retina acquired with a microscope EVOS FL auto 2. The area of 10-20 nuclei per retina was measured and divided to total area of the ONL to calculate the total number of nuclei. The number of photoreceptors in 100 μm per treated/untreated animal was plotted. Mean ± SEM. Two-way ANOVA (* p<0.05, ** p<0.01). N=3.

These data suggest that S1PRs play a critical role in the mechanism of light-induced photoreceptor cell death mediated by rhodopsin M39R. Therefore, blocking S1PRs activity could delay retinal degeneration and reduce light-dependent *Rho^M39R^*phenotypes.

## DISCUSSION

In this study, we developed a new mouse model for sector RP that replicates key features of the disease. The *Rho^M39R/+^* mouse model exhibits mild inferior retinal degeneration by five months of age, replicating both the region-specificity and the late-onset of sector RP (15, 28). Additionally, our data confirmed the critical role of both bright and ambient light in exacerbating the phenotype and accelerating retinal degeneration in sector RP, and highlights that a diagnostic test, such as the ERG, could play a critical role in sector RP progression by inducing further degeneration in the centre of the retina.

C57BL/6J mice are less sensitive to light damage than other strains of mice and particularly albino mice (29); however, C57BL/6J mice carrying the murine *Rho^M39R^*variant showed extreme light sensitivity and retinal degeneration induced by short exposure to bright light via ERG. Some other rhodopsin models show a similar sensitivity to light. For example, short exposure to bright light or fundus photography can cause light-induced toxicity in tg mice carrying the hT17M mouse (30, 31). T17M, together with T4K and T4R, are rhodopsin mutations that inhibit the N-terminal glycosylation of the protein and are associated with sector RP (32-34). A few *Rho^T17M^* KI models have been recently generated, but their sensitivity to light was not studied (35, 36). The role of T4K has been predominantly studied in *X. laevis,* while T4R has been investigated in a naturally occurring dog model. Both models confirmed the critical role of light in rhodopsin T4K and T4R induced retinal degeneration (32-34, 37).

Other rhodopsin amino acid substitutions, such as Y102H and I307N, can lead to light-dependent degeneration after short exposure to bright light (12,000 lux per 5 min) in mutagenized C57BL/6J mouse lines (respectively Tvrm1 and Tvrm4 mice) (38, 39). However, these amino acid changes have not been associated with sector RP in patients. Furthermore, rhodopsin Y102H and I307N did not affect the activity of the retina from early stages, such as M39R or lead to ONL length reduction in mice up to 12 months of age prior to light damage (38). Rhodopsin P23H is also associated with sector RP and its sensitivity to light damage has been investigated in several models including rodents, *X. laevis* and *Drosophila* (30, 40-45). Although *Rho^hP23H^* tg mice are also sensitive to short exposure of bright light, these mice exhibit lower sensitivity compared to *Rho^hT17M^* tg mice (30). In contrast, rhodopsin P23H transgenic rats, which are albino, primarily experience light-induced toxicity following prolonged exposure to bright light (41).

In the T4K *X. laevis* transgenic model, RPE65 deficit can rescue the light-induced toxicity (37). In contrast, in Y102H and I307N mice and P23H *X. laevis*, the ablation of RPE65 exacerbated the phenotype (37, 38). The Moritz’s lab proposed two mechanisms of light-induced degeneration in sector RP: “chromophore-exacerbated” (T4K-like) and “chromophore-mitigated” (P23H-like). The two divergent phenotypes within *RHO* variants associated with sector RP could also depend on the different biological classification: respectively class 4 for T4K and class 2 for P23H (5). *RHO* M39R is a class 4 mutation and similarly to canine rhodopsin T4R shows a faster decay of meta-rhodopsin II, and thermal instability (15, 34). Further investigations will be necessary to elucidate the role of RPE65 and 11-*cis*-retinal in M39R mice.

The *Rho^M39R^* model provides deeper insights into the role of rhodopsin class 4 mutations in OS stability and retina degeneration. Class 4 mutations are hypothesised to cause instability of the OS. The data show that of rhodopsin traffics to the OS in both *Rho^M39R/+^* and *Rho^M39R/M39R^*and the OS has structural alterations. *Rho^M39R/+^* OS show frequent disorganised areas together with a thinner diameter, while rhodopsin levels are comparable to the levels measured in *Rho^+/+^*animals. In heterozygous mice, where 50% of the rhodopsin protein is wild-type RHO, disc formation is similar to the control animals and the presence of rhodopsin M39R causes only a partial OS dysregulation, with vesicular structures intercalated in transversally oriented discs. In contrast, the OS structure changes completely when only rhodopsin M39R is expressed. The intrinsic instability of rhodopsin M39R (15), in combination with the amino acid change occurring in the first transmembrane domain might affect the formation and stability of the discs and therefore lead to highly disorganised and shorter OS. In addition, transcriptomic analysis performed on *Rho^M39R/M39R^* mice reveals the downregulation of *Prph2,* a crucial gene involved in OS disc morphogenesis (46-48).

In 3 week old *Rho^M39R/M39R^*, the amount of rhodopsin is around 70% lower than in *Rho^+/+^* mice. There is also a 30-40% reduction in ONL thickness, which might account for some of this decrease, but does not fully account for the decrease in rhodopsin levels. The reasons why the rhodopsin levels are reduced by more than photoreceptor cell loss could reflect changes in transcription, protein degradation or increased OS turnover. In *Rho^M39R/M39R^* mice, rhodopsin M39R is indeed found in both the ONL and IS. In addition, transcriptomics and immunohistochemistry show *Gfap* increased levels of expression and GFAP-positive Muller glia at 3 weeks of age respectively (data not showed), confirming retinal degeneration in *Rho^M39R/M39R^* mice from an early stage.

Bright light exposure led to a further dysregulation of the disc structure in *Rho^M39R/+^* (TEM data not shown), similar to what was previously reported for other light damage models (49, 50). After light, *Rho^M39R/M39R^* OS become smaller and were mainly formed of vesicular structures. In contrast, rearing the animals in dim red light decreased OS instability in both models and reduce retinal degeneration in *Rho^M39R/M39R^*. In *Rho^M39R/M39R^*, the absence of light facilitates the formation of proper compact discs and improves rhodopsin localization at the OS, reducing the accumulation of the protein in the IS.

Moreover, even though rhodopsin M39R localize at the OS in the presence and absence of WT rhodopsin, there was reduced photoreceptor activity. Rhodopsin M39R, as previously described, can activate phototransduction but with a faster meta-rhodopsin II decay (15). However, the scotopic a wave recorded in *Rho^M39R/+^* at 3 weeks of age was significantly lower than in *Rho^+/+^* mice. In contrast, rearing animals in dim red light rescued the activity in *Rho^M39R/+^,* similar to what has been previously observed in *Rho^P23H/+^* KI mice (43). We hypothesize that after ambient light exposure, rhodopsin M39R might be highly phosphorylated and/or not properly dephosphorylated and that the higher phosphorylation level could lead to increased recovery time and reduced sensitivity to light (51, 52). In dim red light, rhodopsin M39R is unphosphorylated, and therefore it can respond normally to the initial light exposure. Further analysis would be needed to test this hypothesis. However, in *Rho^M39R/M39R^* mice, the scotopic a wave, which is further impaired compared to *Rho^M39R/+^* mice, cannot be rescued in dim red light. Further investigation will be requested to understand if this is due to constitutive instability of the protein and consequently of the OS, despite a measured improvement.

Transcriptomic analysis was performed on the *Rho^M39R/M39R^*, as this model exhibits retinal degeneration under ambient light. These data provided insights into the gene expression changes in a sector RP model in ambient light. Changes in genes of the S1P signalling pathway were identified. The S1P signalling pathway and *de novo* ceramide synthesis are among the pathways that show dysregulation in light damage models (53). In response to light damage, the levels of ceramides and ceramide-related metabolic genes increase in albino rats and Tvrm4 mice (22, 39). In addition, TNF Receptors such as *Tnfrsf1a* respond to changes in sphingolipid levels and have been associated with mechanisms of light-induced toxicity (22-24, 27). Moreover, bright light exposure in albino rodents has been shown to elevate S1P levels, along with the expression of S1P kinases (SphKs) and S1P receptors (S1PRs) (22-24). Nevertheless, mice lacking *SphK2* were still photosensitive to light damage (54). S1P is important for cell proliferation and survival (55). In the retina, it plays a key role in maintaining outer limiting membrane (OLM) stability and regulating cytoskeletal rearrangements in Müller cells (56).

*S1pr3* and *S1pr2* transcripts expression was increased in *Rho^M39R/M39R^* mice compared to *Rho^+/+^* mice. In albino rats, both S1P receptors are expressed in bipolar cells but only S1pr3 localizes in photoreceptors (in both IS and ONL) (23). The S1PRs modulator FTY720 can reduce the light-induced toxicity from bright light in albino rats (22). FTY720 is an approved treatment for Multiple Sclerosis that acts on the central nervous system by modulating inflammation and by improving neuronal functions (57-59). Our data in *Rho^M39/+^* mice confirm the crucial role of FTY720 in preventing light damage by both reducing photoreceptor loss and activation of Müller cells. In *Rho^M39R/M39R^* mice, FTY720 treatment significantly improved photoreceptor survival but did not alter the inflammatory response. In these homozygous mice, Müller cells were already activated by 3 weeks of age under ambient light conditions, even before light-induced damage occurred. Retina activity did not improve with FTY720 treatment; therefore, our findings suggest that FTY720 may function by downregulating the cell death signalling downstream of the rhodopsin M39R response to light.

In conclusion, our data provide new insights into sector RP retinal degeneration and suggest two potential strategies to mitigate light-induced toxicity: reducing light exposure and implementing a therapeutic approach targeting the sphingolipid signalling pathway.

## MATERIALS AND METHODS

### Animal care

All animal procedures were carried out in accordance with UK Home Office regulations under the Animals (Scientific Procedures) Act of 1986 and were approved by the ethics committee of the UCL Institute of Ophthalmology, London, UK. The *Rho^M39R^* line was generated by CRISPR-cas9 HDR technology on C57BL/6J mouse strain. A specific gRNA (Table 1) was used in combination with a donor template containing a change in the spCas9 PAM site (CCA>CTC) and introducing the Met to Arg (ATG>AGG) mutation. *Rho^+/+^* C57BL/6J mice were used as controls. Except for light manipulation experiments, animals were kept on a 12-hour light (20–100 lux) and 12-hour dark (<10 lux) cycle, with free access to food and water. Experiments were conducted using both sexes in equal proportion.

### Drug administration

Mice in dim red light were injected intraperitoneally with vehicle (saline solution) or FTY720 (Bio-techne, Tocris, 1 mg/ml in saline solution). The injections were performed 30 min prior to ERG. FTY720 was injected at a concentration of 10 mg/kg as already described (22). To perform OCT/ERG, mice were anesthetized using Ketamine (Dechra, 1.2 mg/kg) and Medetomidine/Domitor (Orion Pharma, 1 mg/kg); and recovered for longitudinal experiments by administrating Atipamezole/Antisedan (Orion Pharma, 1.3 mg/kg).

### Electroretinogram (ERG)

Mice were dark-adapted overnight and anaesthetised before starting the procedure. Pupils were dilated with topical application of 1% tropicamide (Bausch & Lomb, UK). Electroretinography (ERG) was conducted using a Celeris machine (Diagnosys LLC, UK) under dim red light conditions. Flash stimuli (0.1 ms to 30 ms duration, repetition rate 0.2 Hz) were delivered by the stimulator (log intensity -4 to +3) to assess the scotopic activity of the retina. a- and b-wave responses were recorded, and the data were exported and analysed by Excel (Microsoft) and Prism/GraphPad (Dotmatics).

### Optical coherence tomography (OCT)

Retinae of anaesthetised animals were imaged using the Bioptigen Envis R2300 Spectral-domain ophthalmic imaging system (SDOIS). Images were captured using the InVivoVue Clinic while administrating topical Systane Ultra (Alcon) lubricant eye drops application to prevent cornea dryness. Manual segmentation of the retinal layers was performed using the Bioptigen InVivoVue Diver 2.0 software to measure the outer nuclear layer (ONL) thickness. Exported results were analysed by Excel (Microsoft) and Prism/GraphPad (Dotmatics).

### Transmission electron microscopy (TEM)

Mouse eyes were isolated and were immediately fixed in Karnovsky’s fixative overnight at 4°C. The day after, after rinsing with 0.1 M sodium cacodylate-HCl buffer (pH 7.4), the samples were post-fixed in 1% aqueous osmium tetroxide for 2 hours at room temperature, then dehydrated by passage through ascending alcohols (1 x 30%, 50%, 70%, 90% and 2 x 100%), and two changes of 100% propylene oxide. Samples were infiltrated with 100% resin over 4–6 hours, and then embedded in fresh resin, which was then cured by 48-hour incubation at 60°C. Ultra-thin sections (85 nm) were collected on 150 hex copper TEM grids and were contrasted with Reynold’s lead citrate for 5 minutes at room temperature. Prepared sections were then imaged at TEM JEOL 1400plus operating at 80 kV. Images were captured using a Deben NanoSprint12S camera using the Advanced Microscopy Techniques (AMT) Imaging Software.

### Protein extraction and Western blot

Mouse retinal tissue was lysed in 200 μL RIPA buffer (1% sodium deoxycholate, 150mM NaCl, 1% NP-40, 0.1% SDS, 50mM Tric-HCl pH=7.5) containing 2 % (v/v) protease inhibitor cocktail (PIC; Sigma) and 1% (v/v) Phosphatase inhibitor cocktail (PHIC, Sigma) on ice. Tissue was further disrupted by a brief sonication of 20 s at 0.1 watts using an Ultrasonic Liquid Processor XL-2000 (Misonix Incorporated) and lysates were then centrifuged at 13,000 g for 10 minutes at 4°C for the removal of cellular and DNA debris. Supernatant was then collected, and protein quantification was completed using the Pierce Bicinchoninic acid (BCA) Protein Assay (Thermofisher) following the manufacturers microplate procedure. Absorbance at 562 nm of standards and samples was quantified using a spectrophotometer (Safire, Tecan).

Protein extraction was performed in RIPA Buffer starting from dissected mouse retina. Following protein quantification, 3-5μg of protein of each sample was mixed with Sample buffer 4X (0.025% bromophenol blue, 10% b-mercaptoethanol, 20% glycerol, 5% SDS, 125mM Tris-HCl pH=7). The tissue lysates that were blotted with rhodopsin antibodies remained unheated. The samples were loaded in 10% acrylamide gels and resolved by SDS-polyacrylamide gel electrophoresis (SDS-PAGE). Protein was then transferred to nitrocellulose membranes via wet transfer using Transfer buffer ( 25_mM Tris, pH=8.3, 192_mM glycine, 0.01% SDS and 20% methanol). Membranes were blocked with 5 % (w/v) powdered milk (Marvel) in PBS containing 0.1 % TWEEN-20 (PBS-T, Sigma). Blots were incubated with primary antibodies (Supplementary Table 4), diluted in 2.5-3% milk PBS-T, overnight at 4°C with shaking. The next day, blots were washed three times in PBS-T, followed by a two-hour incubation with HRP-conjugated secondary antibody, diluted in 2.5 % (w/v) milk PBS-T. Following secondary incubation membranes were washed with PBS-T three times and protein detected using Luminata Crescendo or Forte western HRP substrate (Millipore) and imaged with ImageLab on a BioRad ChemiDoc XRS+.

### Immunohistochemistry (IHC)

Enucleated mouse eyes were fixed in a 4% PFA solution overnight, washed with PBS, and incubated in 30% sucrose for 24-48 h, then snap frozen in OCT Embedding matrix (Cell path) and cryosectioned at 10 μm. Slides were washed 3 times with PBS, incubated in Blocking Buffer (BB) (10% serum, 3% BSA in 0.1% triton-X PBS), incubated in primary antibody solution (Table 2), washed 3 times, and finally incubated in secondary antibody solution (Table 2). Each incubation was 1h long at room temperature (RT). A final incubation with 4’,6-diamidino-2-phenylindole (DAPI) in water/PBS for 5min at RT was performed prior to mounting the slides in fluorescence mounting media (DAKO). Images were acquired using a LSM700 laser-scanning confocal microscope (Carl Zeiss) or Stellaris 8 (Leica) and processed using Image J (National Institute of Health, Bathesda, MD, USA) and Illustrator (Adobe).

### Light damage assay

Light damage was induced by repeated ERG every week for 4 weeks starting from mice at 3 or 4 weeks of age. Maximum light intensity was 30000 lux for 2 flashes.

### RNA-sequencing (RNA-seq)

Bulk RNAseq was performed by Azenta. RNA-seq data were analyzed using the R statistical environment (v4.5.0). Raw read counts were processed and normalized with the DESeq2 package (v1.48.0, Bioconductor), applying default settings including size factor normalization and dispersion estimation. Genes with low counts (<10 reads) across all samples were filtered prior to differential expression testing. Differentially expressed genes (DEGs) were identified using the DESeq2 Wald test, with significance thresholds set at adjusted *p*-value < 0.05 following Benjamini–Hochberg correction for multiple testing. Volcano plots were generated using the ggplot2 package (v3.5.2) and ggrepel (v0.9.6) for label repelling, with genes color-coded according to significance thresholds. Heatmaps were constructed using the pheatmap package (v1.0.12). Variance-stabilizing transformation (VST) was applied to normalized counts using DESeq2, and average expression values per condition were calculated. Expression values were centered by subtracting the global mean across all genes and conditions. KEGG pathway enrichment analysis was conducted using the clusterProfiler package (v4.16.0) after converting gene symbols to Entrez IDs via org.Mm.eg.db (v3.21.0). Enrichment was performed against the Mus musculus KEGG database (*organism* = "mmu"). The top 20 enriched KEGG pathways were visualized using the enrichplot package (v1.28.0).

## ACKNOWLODGMENTS

These authors gratefully acknowledge Hannah Poultney for the help in the animal care and Prof. Robert Molday (University of British Columbia) for the gift of the 1D4 antibody.

## AUTHOR CONTRIBUTIONS

Conceptualization, R.G., M.E.C. Formal analysis, R.G., K.Z.; Funding Acquisition, M.E.C.; Investigation, R.G., K.Z; Methodology, R.G., K.Z., K. H., D.P., S.E.N, A.B., R.A., M.A., D.A., D. S. S., Y.L.; Project Administration, M.E.C.; Resources, M.E.C..; Supervision, M.E.C..; Validation R.G., K.Z; Visualization; R.G., K.Z.; Writing – original draft; R.G., K.Z, M.E.C.; Writing – review and editing, R.G., K.Z., K. H., D.P., S.E.N, A.B., R.A., M.A., D.A., D. S. S., Y.L, R.C., M.E.C.

