## Supplementary material for "Preventing light-induced toxicity in a new mouse model of rhodopsin sector retinitis pigmentosa": Tables and Supplementary Figures

**Supplemental Table 1**

| **gRNA** | GAACATGTACGCTGCCAGCATGG |
| --- | --- |
| **donor template** | GTGGTGCGGAGCCCCTTCGAGCAGCCGCAGTACTACCTGGCG  GAACCATGGCAGTTCTC**t**A**g**GCTGGCAGCGTACATGTTCCTGCT  CATCGTGCTGGGCTTCCCCATCAACTTCCTCACGCTC |

ATG (Met) -> A**g**G (Arg)

Change NGG to NAG (PAM site) TCC ->TC**t**

**Supplemental Table 2**

| **Protein effect** | **Nucleotide change** | **Location** | **Sanger Sequencing** |
| --- | --- | --- | --- |
| Met39Arg | c.116T>G | TM helix I | 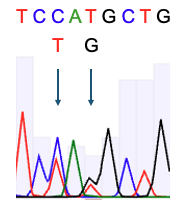 |

**Supplemental Table 3**

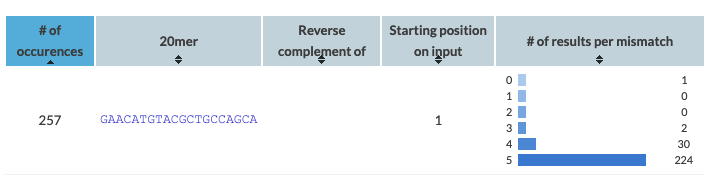

In silico prediction of gRNA Off-target occurrences using Off-Spotter (<https://cm.jefferson.edu/Off-Spotter/>). *Rho* was the only gene listed with 0 mismatch.

**Supplemental Table 4**

| **Antibody** | **Host** | **Supplier** | **Working concentration** |
| --- | --- | --- | --- |
| anti-rhodopsin, Rho-4D2 | Mouse | Millipore, MABN15 | WB 1:1000  IHC 1:1000 |
| anti-rhodopsin, Rho-1D4 | Mouse | Professor Molday, UBC | 1:1000 |
| Anti-GFAP | Rabbit | Dako, GA524 | 1:500 |
| Alex Fluor 488 or 555 | Donkey anti-mouse or rabbit | Thermo Fisher Scientific | 1:1000 |

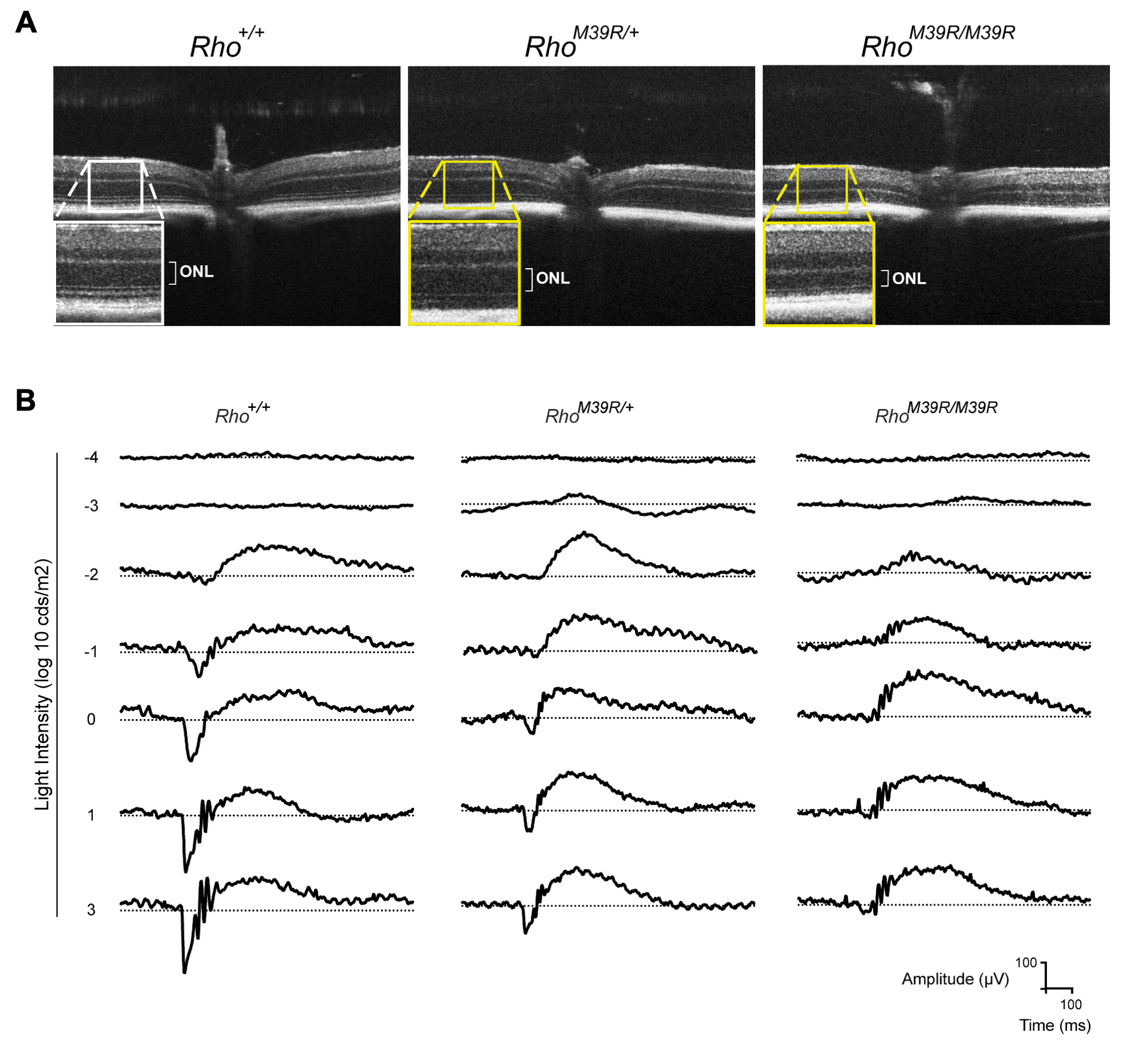

**Supplementary Figure 1.** (**A**) Representative images of 3 weeks old *Rho^+/+^*, *Rho^M39R/+^* KI and *Rho^M39R/M39R^* KI mouse retina acquired by OCT using Bioptigen SD-OCT. The inset shows the ONL at higher magnification. ONL = Outer Nuclear Layer. (**B**) ERG recorded at different light intensities using a Celeris ERG system (Diagnosys LLC).

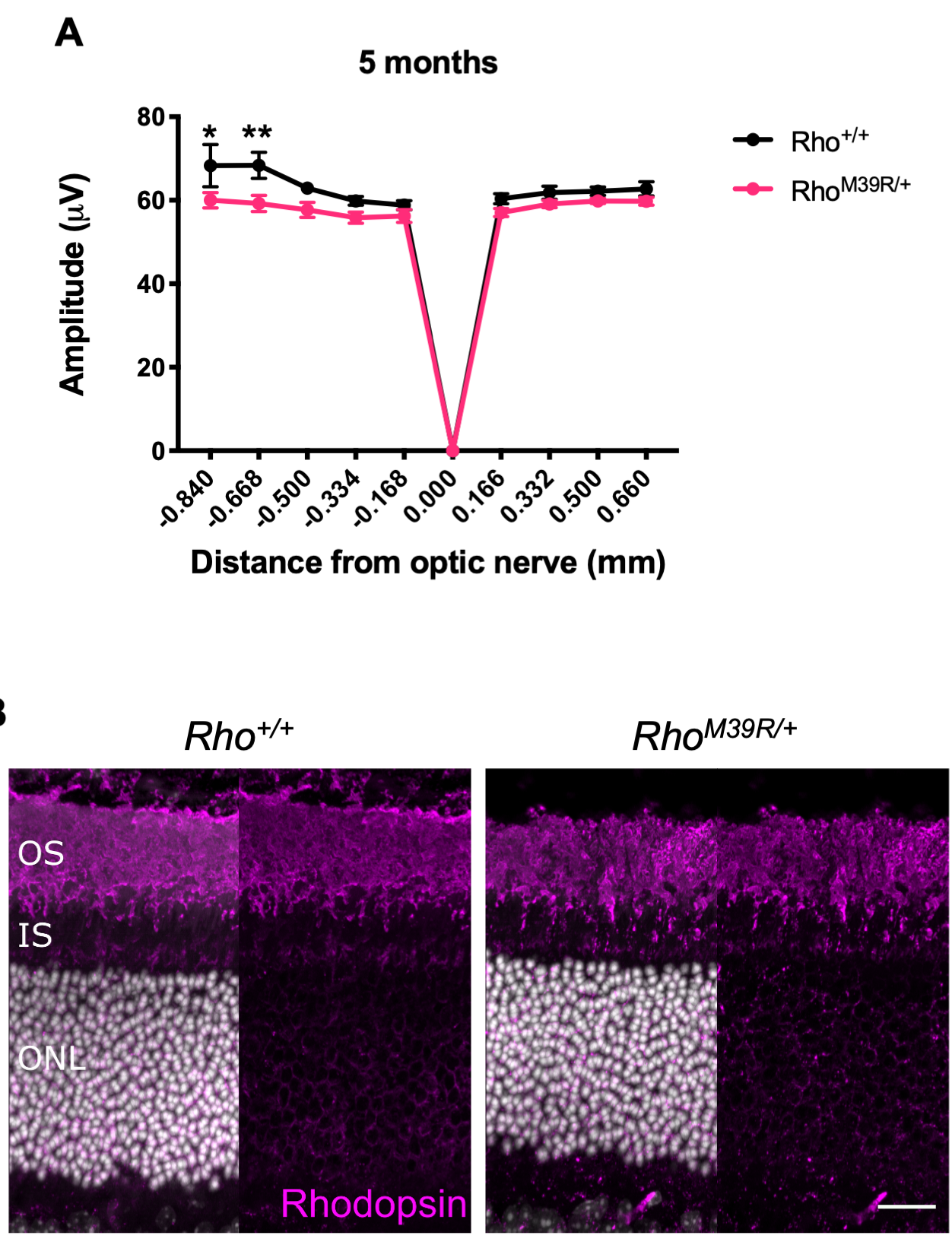

**Supplementary Figure 2.** (**A**) ONL thickness was measured by OCT in the central retina of *Rho^+/+^* and *Rho^M39R/+^* KI at 5 months. Mean ± SEM. Two-way ANOVA. Sidak’s multiple comparisons test (* p<0.05, ** p<0.01). N=3/4. (**B**) Rodent eyes were fixed in 4% PFA, incubated in 30% sucrose for 1-2 days, embedded in OCT (embedding matrix), cryosectioned and stained with DAPI and anti-rhodopsin-1D4 antibody. Scale bar=20μm.

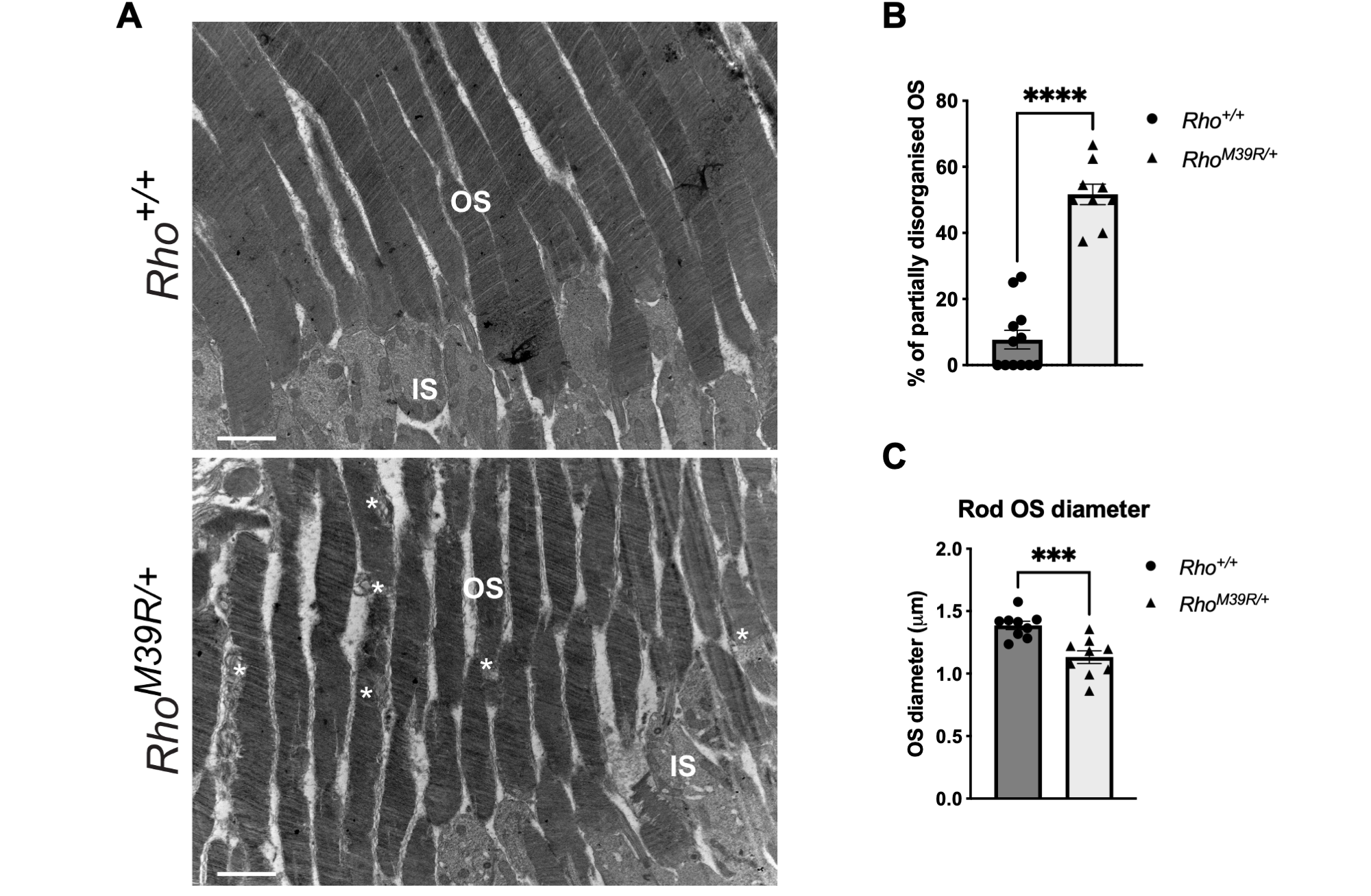

**Supplementary Figure 3.** (**A**) Representative images of *Rho^+/+^* and *Rho^M39R/+^* KI mouse retina acquired by TEM. Stars indicates disorganised areas along the RO structure. OS = Outer Segment, IS = Inner Segment. Scale bar=2μm. (**B**) The percentage of partially disorganised RO in *Rho^+/+^* and *Rho^M39R/+^* KI mouse retina was measured in TEM pictures and plotted. Mean ± SEM. Mann-Whitney test. 9 pictures from 2 independent animals. (**C**) The diameter of the OS was measured in *Rho^+/+^* and *Rho^M39R/+^* KI mouse retina. Mean ± SEM. Mann-Whitney test. 12-9 pictures from 2 independent animals.

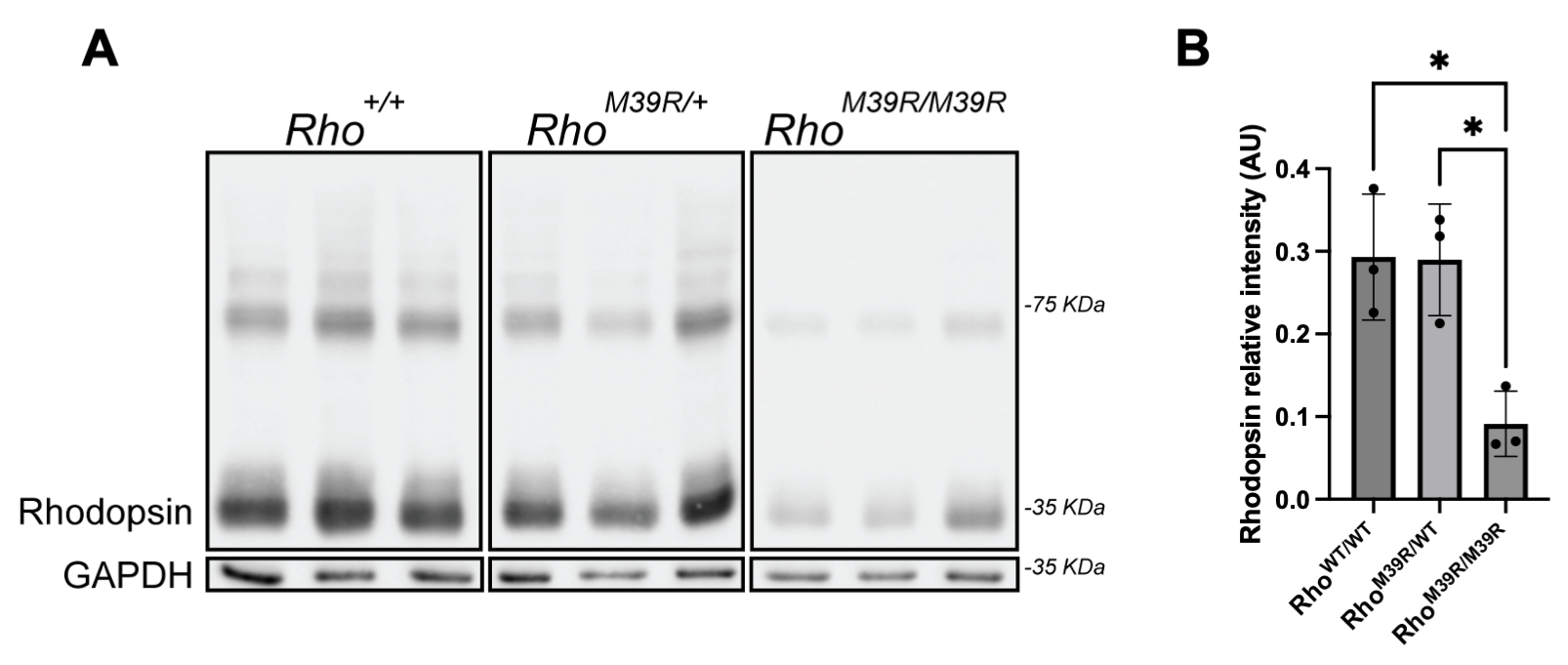

**Supplementary Figure 4** (**A**) Immunoblot of 3 weeks old *Rho^+/+^*, *Rho^M39R/+^* and *Rho^M39R/M39R^* KI mouse retina stained with rhodopsin-4D2 antibody. Anti-GAPDH was used as a loading control. (**B**) Plot of rhodopsin-4D2 relative intensity (34kDa band) measured using Fiji. Values were normalised on GAPDH intensity signal. Mean ± SD. One-way ANOVA. (* p<0.05).

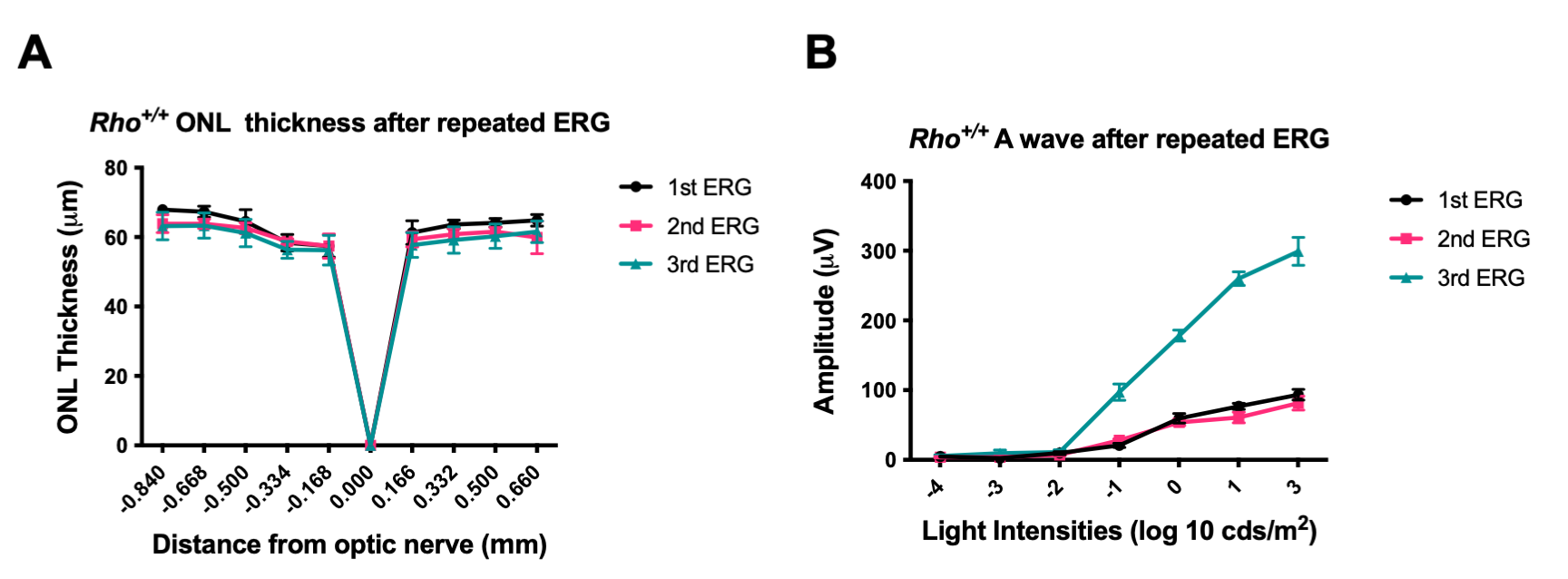

**Supplementary Figure 5.** (**A**) ONL thickness of *Rho^+/+^* after 3 different rounds of ERG. The values were measured by OCT. Mean ± SEM. Two-way ANOVA. N=4 (**B**) The scotopic A was also measured in *Rho^+/+^*. Mean ± SEM. Two-way ANOVA. N=4.

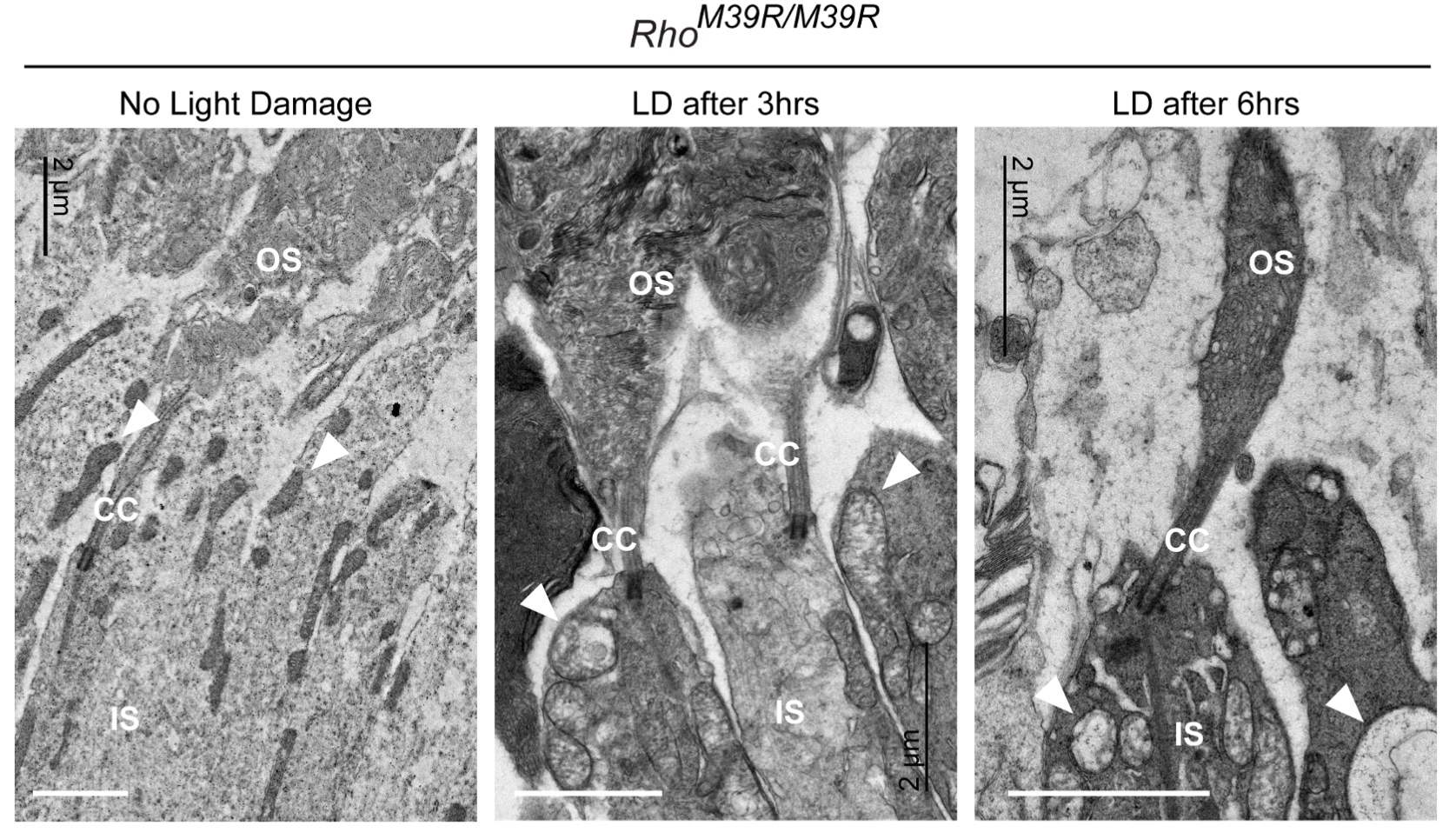

**Supplementary Figure 6.** TEM of *Rho^M39R/M39R^* KI mouse showing the ultrastructure of both OS and IS. Arrows indicate mitochondria. OS = Outer Segment, CC = connecting cilium, IS = Inner Segment. Scale bar=2μm.

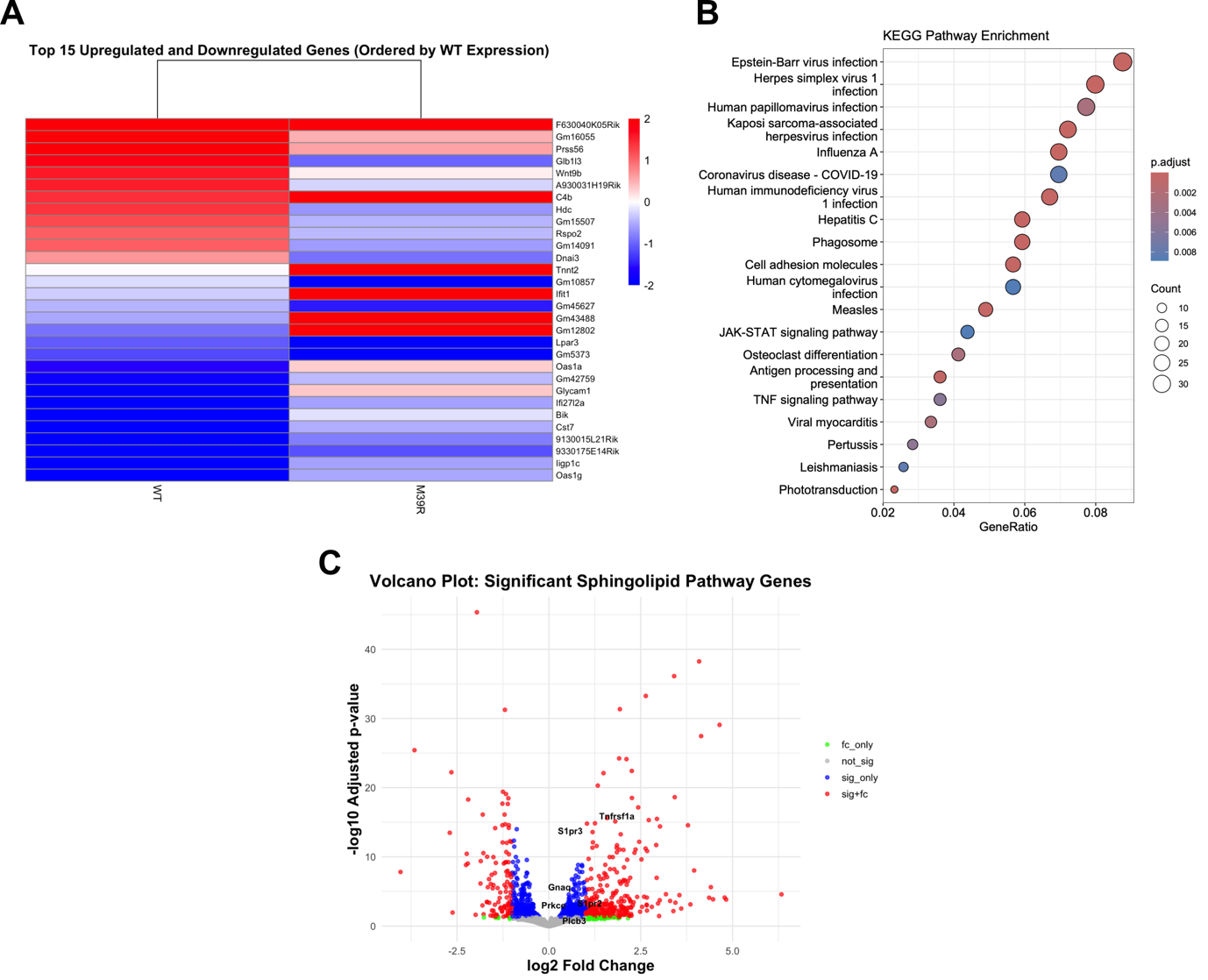

**Supplementary Figure 7. Transcriptomic analysis of** *Rho^M39R/M39R^* KI mouse compared to *Rho****^+/+^*.** (**A**) Heatmap of the top 15 upregulated and downregulated genes in *Rho^M39R/M39R^* KI mouse compared to *Rho^+/+^*. (**B**) KEGG enrichment pathway analysis. (**C**) Volcano plot of sphingolipids signalling pathway genes upregulated in our cohort. fc_only = only fold change; not_sign = not significant; sig_only = only significant; sig+fc = significant plus fold change.

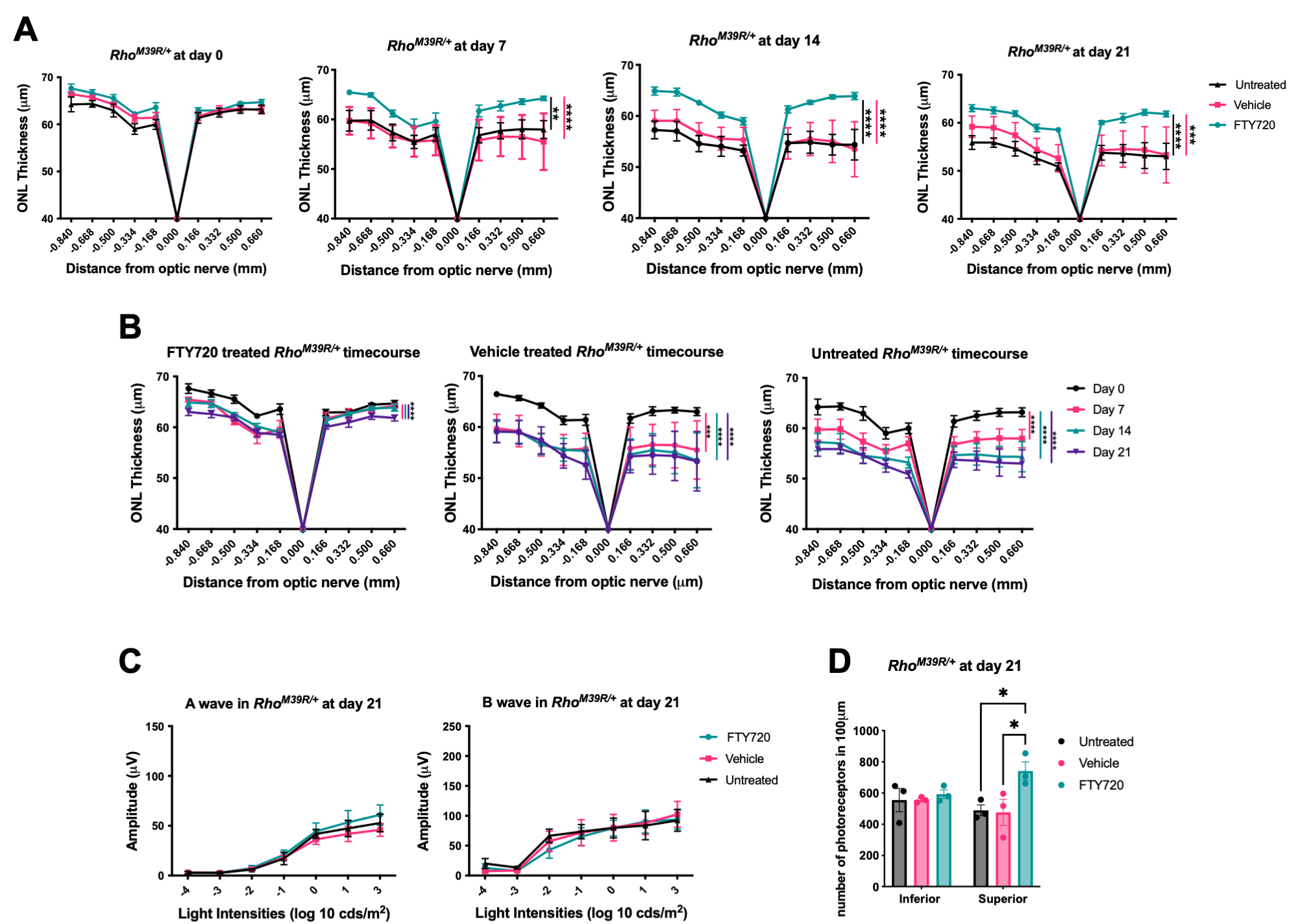

**Supplementary Figure 8.** (**A**) ONL thickness of *Rho^M39R/+^* KI mouse retina at each round of ERG: day 0, day 7, day 14 and day 21. Dark-adapted mice were intraperitoneally injected with 10 mg/Kg of FTY20 or with vehicle (saline solution) 30 min prior to the ERG. Untreated mice were also analysed. Mean ± SEM. Two-way ANOVA. (**** p<0.0001, *** p<0.001, ** p<0.01). N=5/7 (**B**) ONL thickness of FTY20-treated, vehicle-treated and untreated *Rho^M39R/+^* KI mice measured over time. Mean ± SEM. Mixed-effect analysis (**** p<0.0001, *** p<0.001). N=5/7. (**C**) A and B wave of FTY20-treated, vehicle-treated and untreated *Rho^M39R/+^* KI mice measured at the end of the light damage assay (day 21). Mean ± SEM. N=3/4. (**D**) IHC of untreated, vehicle or FTY70 treated *Rho^M39R/M39R^* superior retina after light damage. The cryosections were stained with DAPI. The number of photoreceptors in the ONL was measured at 200 to 400 μm from the optic nerve in the inferior and superior retina. The analysis was performed on images of the central retina acquired with a microscope EVOS FL auto 2. The area of 10-20 nuclei per retina was measured and divided to total area of the ONL to calculate the total number of nuclei. The number of photoreceptors in 100 μm per treated/untreated animal was plotted. Mean ± SEM. Two-way ANOVA (* p<0.05, ** p<0.01). N=3. Two-way ANOVA (* p<0.05). N=3.

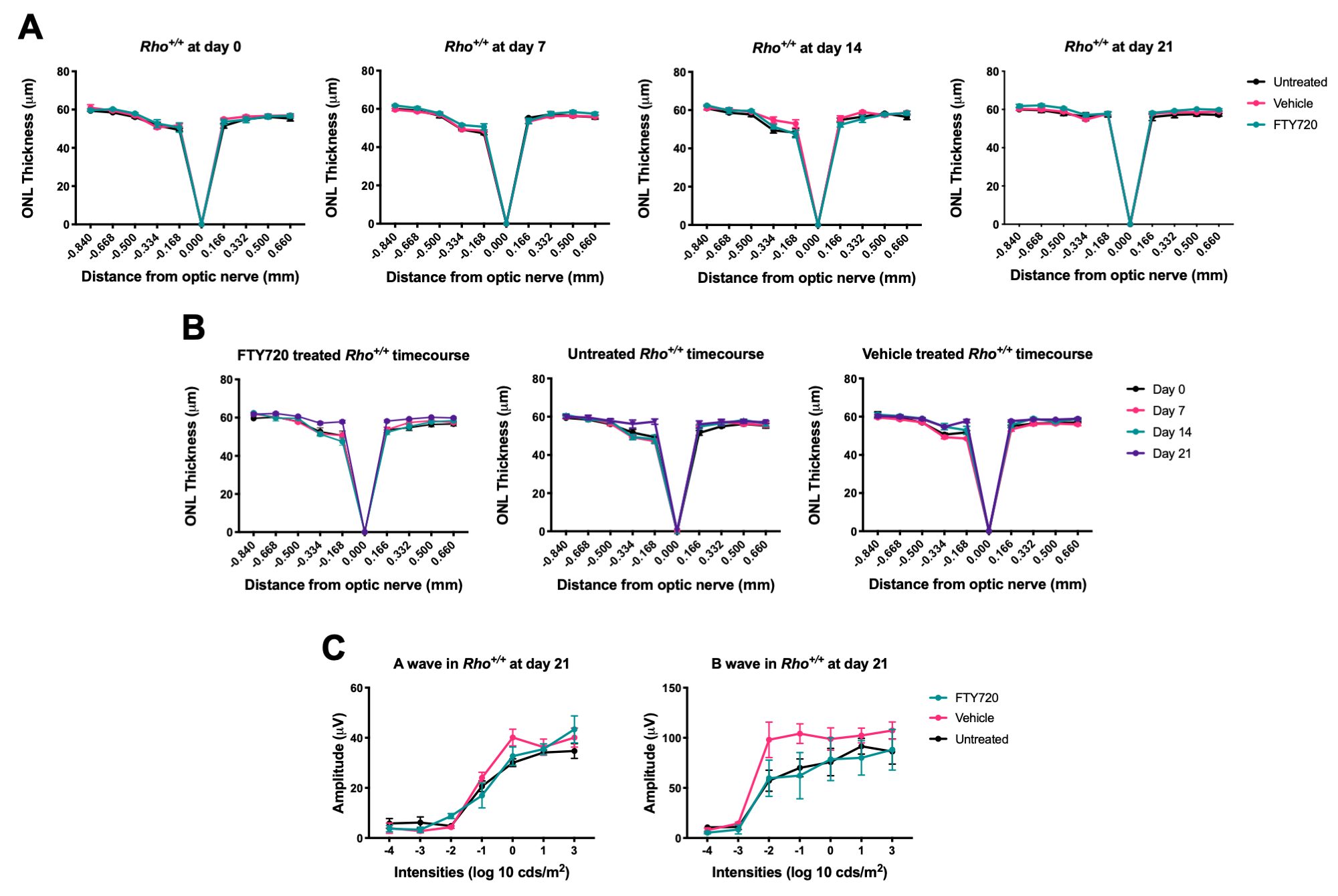

**Supplementary Figure 9.** (**A**) ONL thickness of *Rho^+/+^* KI mouse retina at each round of ERG: day 0, day 7, day 14 and day 21. The measurements were taken by OCT. Dark-adapted mice were intraperitoneally injected with 10 mg/Kg of FTY20 or with vehicle (saline solution) 30 min prior to the ERG. Untreated mice were also analysed. (**B**) ONL thickness of FTY20-treated, vehicle-treated and untreated *Rho^+/+^* KI mice measured over time. (**C**) Scotopic A and B wave was also calculated. Mean ± SEM. Two-way ANOVA. N= 3/4

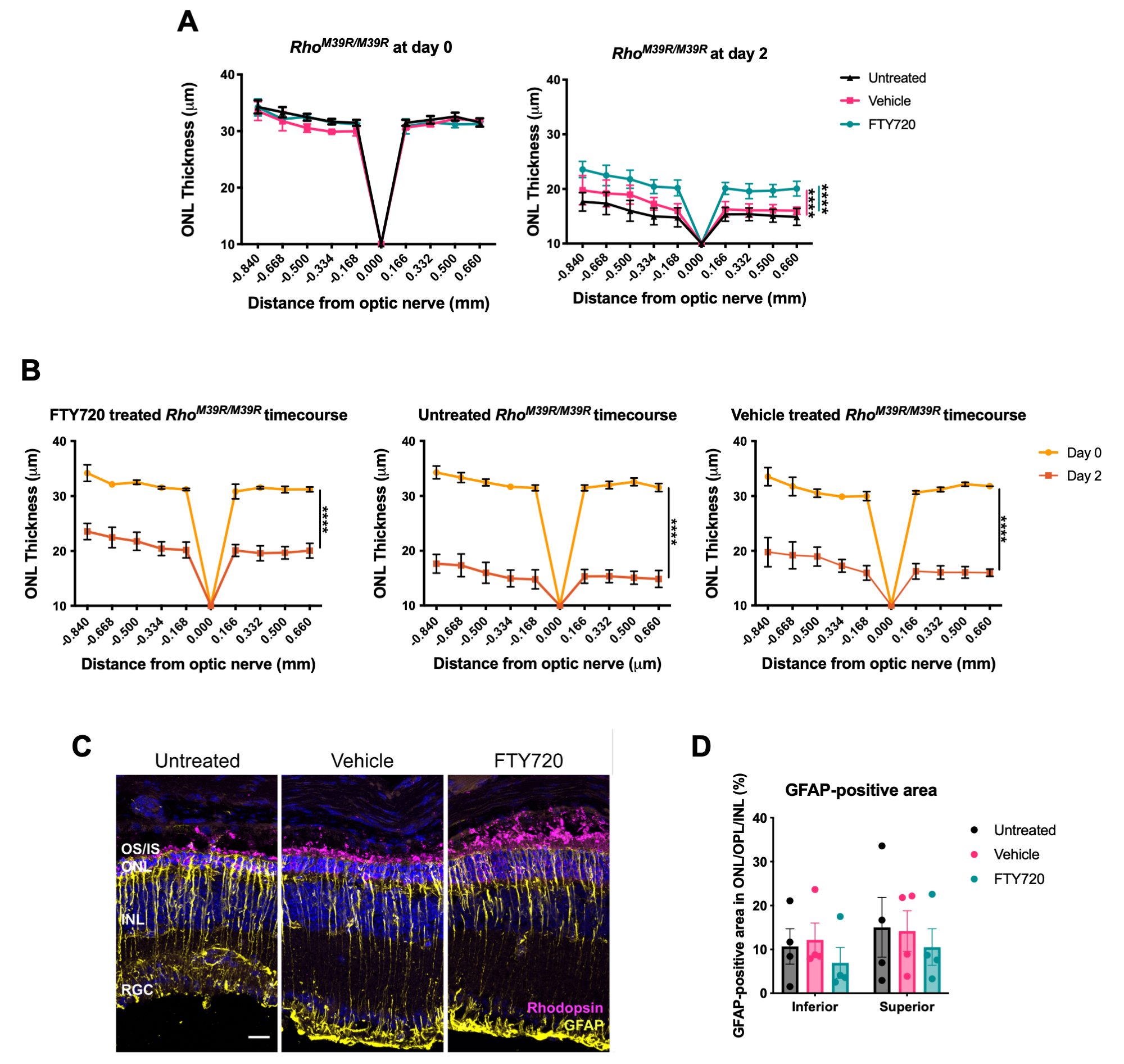

**Supplementary Figure 10.** (**A**) ONL thickness of *Rho^M39R/M39R^* KI mouse retina at day 0 and day 2, after a single ERG. Measurements were taken by OCT. Dark-adapted mice were intraperitoneally injected with 10 mg/Kg of FTY20 or with vehicle (saline solution) 30 min prior to the ERG. Mean ± SEM. Two-way ANOVA. (**** p<0.0001) N=5/6. (**B**) ONL thickness of FTY20-treated, vehicle-treated and untreated *Rho^M39R/M39R^* KI mice measured over time. Mean ± SEM. Mixed-effect analysis (**** p<0.0001). N=5/7. (**C**) IHC of untreated, vehicle or FTY70 treated *Rho^M39R/M39R^* superior retina after light damage. The cryosections were stained with DAPI, anti-rhodopsin-4D2 (in magenta) and anti-GFAP (in yellow). Scale bar=20μm. (**D**) The % of GFAP-positive area in the ONL, OPL and INL was also measured. Mean ± SEM. Two-way ANOVA. N=4.

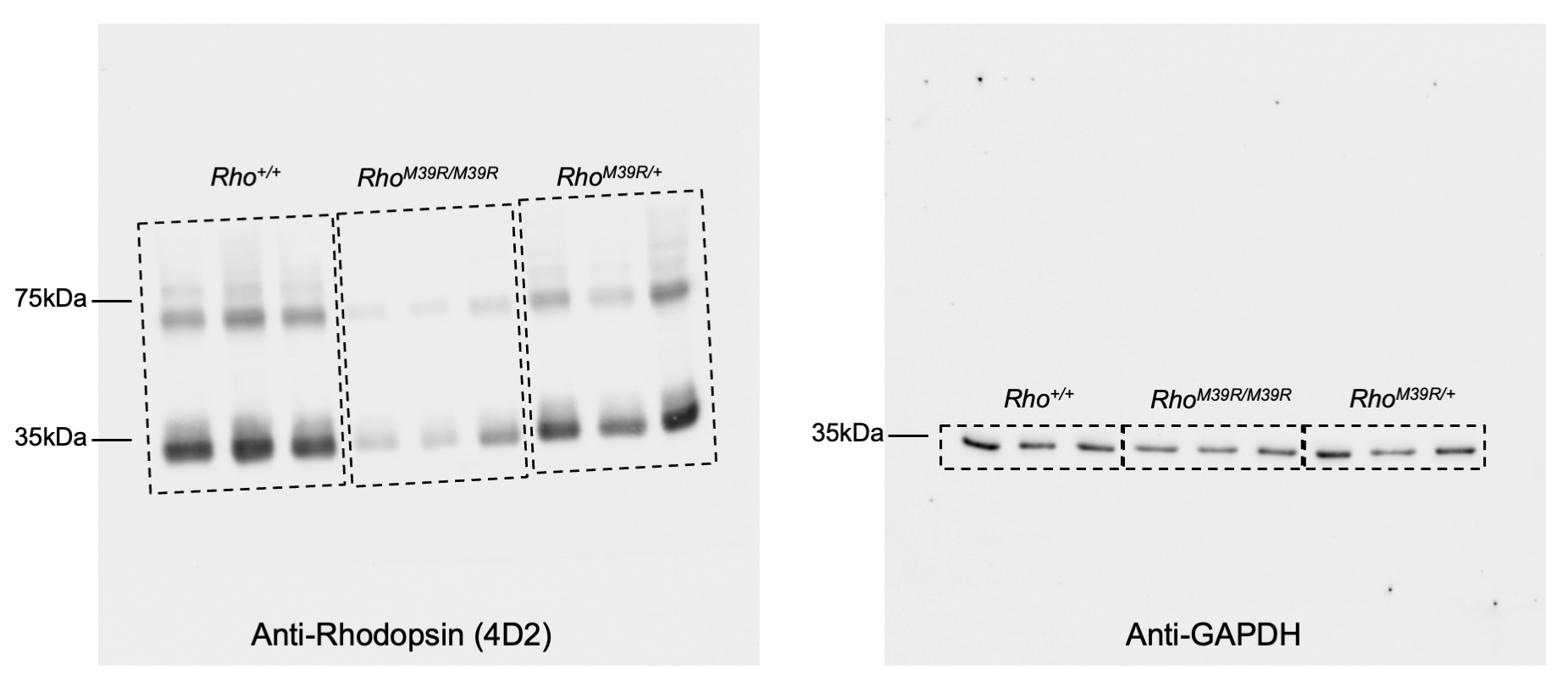

**Supplementary Figure 11.** Uncropped western blot images.
